# Dissection of centrosomal γ-TuRC activation pathways controlling microtubule density in interphase cells

**DOI:** 10.64898/2026.08.17.745143

**Authors:** Yinlong Song, Dipti Rai, Lilian M. Sluimer, Marèl F.M. Spoelstra, Quinten J. Kleijnen, Bert Jan Korte, Silas T. Koot, Kelly E. Stecker, Fangrui Chen, Anna Akhmanova

**Author notes:** Equal contribution (Yinlong Song, Dipti Rai).

## Abstract

Animal microtubule-organizing centers, including the centrosome and the Golgi apparatus, regulate microtubule nucleation and anchoring through the γ-tubulin ring complex (γ-TuRC) and CAMSAP-mediated minus-end stabilization. However, functional redundancy between these pathways has impeded dissection of their contributions to controlling microtubule organization and density. Here, we addressed this problem using combinatorial gene knockouts, protein depletions and Expansion Microscopy. By simultaneously eliminating CAMSAP2 and the γ-TuRC-targeting proteins AKAP450, pericentrin, CDK5RAP2, myomegalin, ninein and AKNA, we generated viable RPE1 cells that lack both Golgi-derived microtubules and γ-TuRC localization within the pericentriolar material and at subdistal appendages. Despite the disruption of these major microtubule-organizing pathways, overall microtubule density was only partially reduced. The remaining microtubules depended on CEP192 and NEDD1, which, together with ch-TOG, can activate γ-TuRC at the centriole wall, in acentriolar cells, and in biochemical reconstitution assays. Our results demonstrate that in the absence of CAMSAP-mediated stabilization, interphase microtubule formation strongly relies on γ-TuRC activation, which occurs through several redundant pathways.

## Introduction

Microtubules (MTs) are cytoskeletal filaments, which serve as tracks for intracellular transport and a scaffold for positioning of organelles and control of cell morphology, polarity and signaling. All these functions critically depend on MT density, which can vary from a few dozen MTs in immune cells to several thousand in the pillar cells of the cochlea (reviewed in ^1^). MT density is determined by the collective action of multiple proteins, but a systematic view of this process is lacking. For example, it is unclear to which extent the first step of MT formation, nucleation, can control MT density. Cellular MTs are typically nucleated by the γ-tubulin ring complex (γ-TuRC), a cone-shaped structure that serves as a template for MT assembly and caps MT minus ends (reviewed in ^2–4^). In vitro work demonstrated that by itself, γ-TuRC has low activity, and it undergoes a conformational change during MT nucleation ^5–11^. The best-studied activator of γ-TuRC is CDK5RAP2 ^11–16^, though it is not an essential protein ^17–21^. Two other known γ-TuRC activators are the MT polymerase XMAP215/ch-TOG and MT growth regulators CLASP1 and 2 (CLASPs) ^22–24^, which can also promote MT nucleation independently of γ-TuRC ^25–27^.

γ-TuRC activators are typically concentrated at MT-organizing centers (MTOCs) such as the centrosome – an organelle that consists of two centrioles surrounded by pericentriolar material (PCM) (reviewed in ^28–30^. Major PCM components are γ-TuRC adaptors ^3, 31^, including pericentrin (PCNT) ^32–34^ and AKAP450; the latter also serves as an essential γ-TuRC-targeting factor on Golgi membranes ^35^. Both PCNT and AKAP450 can recruit CDK5RAP2 and its paralog myomegalin (MMG) ^17, 36^. Another, poorly studied potential γ-TuRC-targeting protein is AKNA, which localizes to the subdistal appendages of the mother centriole and regulates centrosomal MT formation during neuronal development ^37^. Ninein (NIN), another component of subdistal appendages, was also implicated in γ-TuRC recruitment and MT organization ^38, 39^.

A different type of γ-TuRC binding partner is the WD40 repeat-containing protein NEDD1 ^15, 40–42^. NEDD1 interacts with the centrosomal adaptor CEP192 ^43, 44^, which plays a role in centrosomal MT organization ^18, 43–46^, is required for centriole duplication ^47, 48^ and the formation of the mitotic spindle ^44, 45, 49–51^. During mitosis, CEP192 participates in MT nucleation in a complex with XMAP215/ch-TOG ^52^. Based on experiments in *Xenopus* egg extracts and mammalian cells, this process depends on CEP192-mediated recruitment of NEDD1, γ-TuRC and two mitotic kinases, Plk1 and Aurora A, which phosphorylate and activate different components of the complex ^49, 52–56^.

Since XMAP215/ch-TOG and its counterparts in different systems bind free tubulin, it has been proposed that the major function of the centrosome is not to activate γ-TuRC but simply to form a condensate that concentrates tubulin dimers ^57, 58^. Although popular, this notion has not been critically tested in cells, because while many loss-of-function studies of individual centrosome components have been performed, only a few studies attempted to systematically dissect the roles of these proteins in the same cellular model ^17, 18, 49, 59, 60^. Importantly, all studied perturbations of PCM proteins had an effect on mitosis, when the centrosome activity is upregulated ^28, 61^, but had only a limited impact on the overall MT abundance in interphase cells. This is because in addition to centrosomes, interphase mammalian cells can also nucleate MTs from the Golgi complex ^62^, and non-centrosomal MT minus ends can be efficiently stabilized by CAMSAPs ^63^. Centrosome loss or impairment can even increase interphase MT density, by promoting MT release from the nucleation sites and their stabilization by CAMSAPs ^18, 23^.

Here, we systematically dissected centrosomal pathways of MT formation in interphase cells, using RPE1 cells that lack AKAP450 and CAMSAP2, thereby eliminating Golgi-dependent MT nucleation and minus-end stabilization ^17^. In these cells, most MTs originate from the centrosome, and we used combinatorial knockouts and depletion of different centrosomal γ-TuRC-binding and activating proteins to determine their relative contributions to γ-TuRC recruitment to different centrosomal sites, MT nucleation and control of MT density. Our data show that in the absence of CAMSAPs, loss of γ-TuRC cofactors leads to a profound reduction in MT density, demonstrating that interphase centrosomal MT formation requires γ-TuRC activation, and not just concentration of soluble tubulin.

## Results

### Simultaneous loss of major PCM proteins has a modest effect on interphase MT density

To probe the functional redundancy between different PCM proteins, we used a series of knockout (KO) hTERT-RPE1 cells (RPE1 hereafter). The commonly used RPE1 cells are puromycin-resistant, which can be inconvenient for genome editing, but this problem was solved in the Holland lab by knocking out the puromycin acetyltransferase gene ^64^. We observed no differences in MT organization between the original and puromycin-sensitive RPE1 cells. Some of the cell lines we used, described in our previous studies ^17, 65^, were generated in puromycin-resistant RPE1 background. Several new cell lines described in this study were generated using puromycin-sensitive cells (Fig. S1a-d, Table S1). Our cell line collection included knockouts of genes encoding p53, CAMSAP2, AKAP450, CDK5RAP2, MMG (both MMG1 and MMG8 isoforms), PCNT, NIN and AKNA, in different combinations (Fig. S1a-e, Fig. 1a). As described previously ^17^, cells lacking CAMSAP2, AKAP450, CDK5RAP2 and MMG were viable, while PCNT knockout cells could proliferate only on the background of p53 knockout, likely due to p53-dependent cell cycle arrest triggered by prolonged mitosis in the absence of PCNT ^65^. We note that the AKAP450 guide RNAs used in this study do not target the short C-terminal AKAP450 isoform (NP_001366206), which can be detected at the centrosome using antibodies against AKAP450 C-terminus ^18^. Similarly, a short CDK5RAP2 isoform (AAH04526), lacking the γ-TuRC-binding CM1 domain, could be expressed in CDK5RAP2 knockout cell lines.

**Figure 1.**
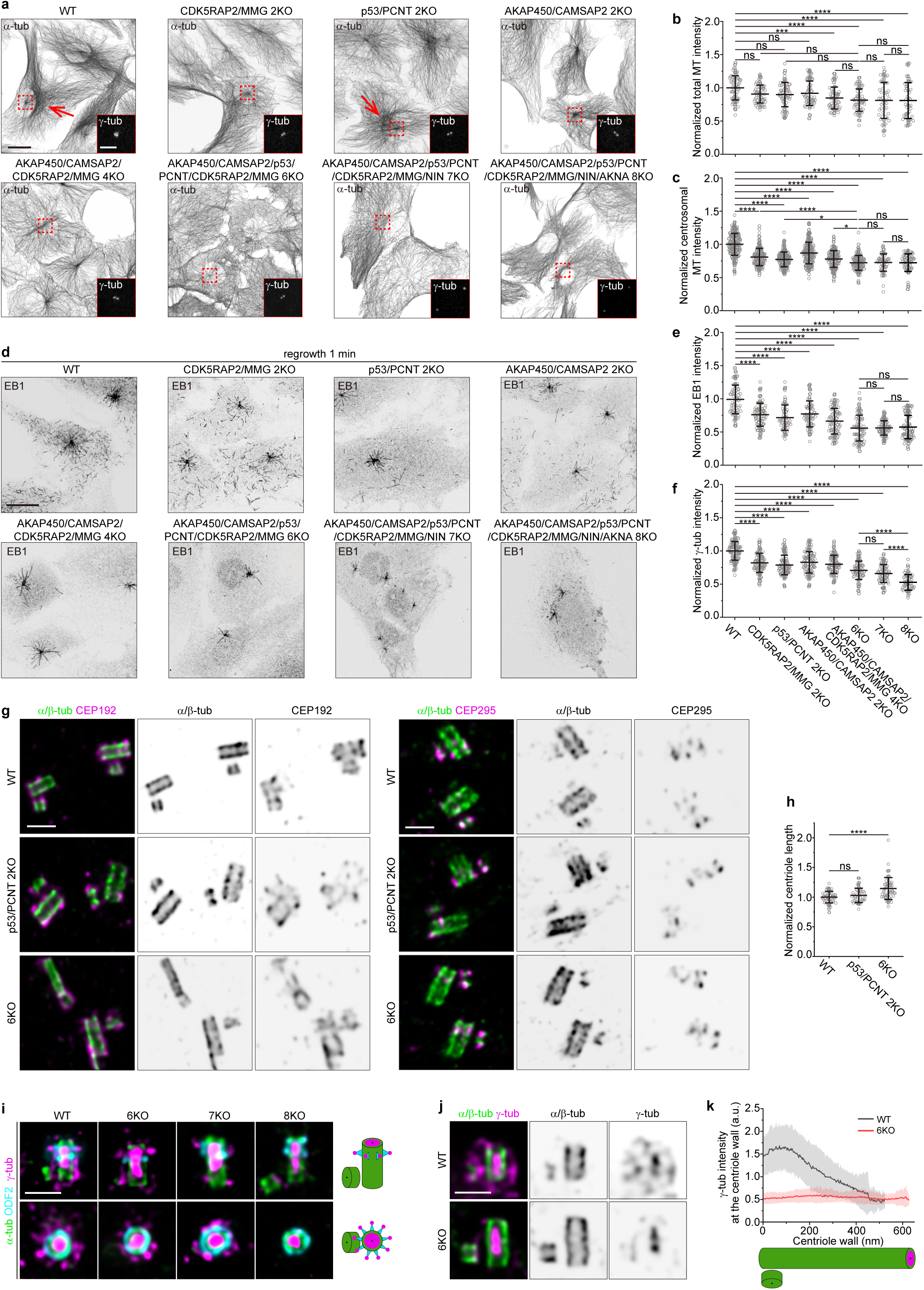
Loss of PCM components selectively impairs centrosomal γ-TuRC pools and reduces interphase microtubule density. **a,** Representative immunofluorescence images of the indicated RPE1 cell lines stained for α-tubulin (grey). Insets show higher-magnification views of boxed regions, with γ-tubulin signal in white. Red arrows indicate Golgi MTs. Scale bar: 20 µm (main image); 5 µm (inset). **b, c, f,** Quantification of total MT intensity (**b**) per cell (mean ± s.d.), centrosomal MT intensity (**c**) per cell (mean ± s.d.) and centrosomal γ-tubulin intensity (**f**) per centrosome (mean ± s.d.) from the experiments shown in **a**. Values are normalized to the wild type average. The numbers of cells analyzed for each cell line (in the same order as the graph) are as follows: for total MT (**b**): 63, 60, 62, 60, 59, 58, 49 and 44 (from four independent experiments); for centrosomal MT (**c**): 190, 146, 170, 192, 160, 130, 58 and 62 (from three independent experiments); for centrosomal γ-tubulin (**f**): 107, 96, 97, 96, 102, 75, 76 and 54 (from three independent experiments). Statistical significance was determined using one-way ANOVA with Tukey’s post hoc test for multiple comparisons (*0.01 < *P* < 0.05; \*\*\**P* = 0.0004; \*\*\*\**P* < 0.0001; ns, not significant, *P* > 0.05). **d,** Representative immunofluorescence images of the indicated RPE1 cell lines showing MT regrowth 1 min after nocodazole washout. Cells were stained for EB1 (grey). Scale bar: 20 µm. **e,** Quantification of centrosomal EB1 intensity per cell (mean ± s.d.) from the experiments shown in **d**. Values are normalized to the wild type average. The numbers of cells analyzed for each cell line (in the same order as the graph) are 75, 84, 63, 71, 83, 75, 97 and 78, from three independent experiments. Statistical significance was determined using one-way ANOVA with Tukey’s post hoc test for multiple comparisons (\*\*\*\**P* < 0.0001; ns, not significant, *P* > 0.05). **g,** Representative U-ExM images of centrioles from RPE1 wild type, p53/PCNT 2KO and 6KO cell lines, stained for α- and β-tubulin (green), CEP192 (magenta) and CEP295 (magenta). Scale bar: 0.5 µm. **h,** Quantification of centriole length (mean ± s.d.) from the experiments shown in **g**. Values are normalized to the wild type average. The numbers of cells analyzed for each cell line (in the same order as the graph) are 53, 45 and 58, from two independent experiments. Statistical significance was determined using one-way ANOVA with Tukey’s post hoc test for multiple comparisons (\*\*\*\**P* < 0.0001; ns, not significant, *P* = 0.5835). **i,** Representative U-ExM images of centrioles from RPE1 wild type, 6KO, 7KO and 8KO cell lines, stained for α- and β-tubulin (green), ODF2 (cyan) and γ-tubulin (magenta). Scale bar: 0.5 µm. **j,** Representative U-ExM images of centrioles from RPE1 wild type and 6KO cell lines, stained for α- and β-tubulin (green) and γ-tubulin (magenta). Scale bar: 0.5 µm. **k,** Quantification of γ-tubulin intensity (mean ± s.d.) along the centriole from the proximal to the distal end (shown in **j**) in RPE1 wild type (black, n = 28) and 6KO (red, n = 31) cell lines. n indicates the number of centrioles analyzed, from two independent experiments.

While none of the knockouts affected the expression of α- or γ-tubulin (Fig. S1b-d), we observed some reduction of total interphase MT density in all studied knockout cell lines (Fig. 1b). As expected, wild type (WT) and p53/PCNT double knockout (2KO) cells had clear MT enrichment on one side of the cell nucleus (Fig. 1a, red arrows), corresponding to Golgi MTs, which depend on AKAP450, CAMSAP2, CDK5RAP2 and MMG ^17^. This MT population was reduced in CDK5RAP2/MMG 2KO and lost in all cell lines lacking AKAP450 and CAMSAP2 ^17^, whereas the centrosomal MT aster persisted (Fig. 1a). Quantification of the centrosome-associated MT population showed that simultaneous knockout of CAMSAP2 and AKAP450 had the mildest effect, as expected ^17^ (Fig. 1c). The reduction of centrosomal MTs in CDK5RAP2/MMG 2KO (~20%) and AKAP450/CAMSAP2/CDK5RAP2/MMG quadruple knockout (4KO) cells (~23%) was quantitatively similar to that of cells lacking PCNT (~23%), whereas 6KO cells, lacking p53, AKAP450, CAMSAP2, CDK5RAP2, MMG and PCNT displayed a somewhat stronger (~28%) loss of pericentrosomal MT density (Fig. 1a,c). Knocking out NIN in 6KO cells (7KO, Fig. S1c) induced centrosome separation due to the abrogation of the centrosome linker, as described previously ^66, 67^, but did not further reduce centrosomal MT density when the signals from both asters were summed (Fig. 1a,c). The same was true for cells where AKNA, a subdistal appendage protein present at the mother but not the daughter centriole ^37^, was also knocked out (8KO, Fig. 1a,c and Fig. S1d,e). The reduction of total MT intensity was also similar (~19%) in 6KO, 7KO and 8KO cells. These data indicate that PCM proteins PCNT, AKAP450, CDK5RAP2 and MMG redundantly control a part of the centrosomal MT population, whereas the contribution of centriolar appendage proteins NIN and AKNA to MT density in RPE1 cells is minor.

This conclusion was supported by nocodazole washout assays, which assess the ability of MTOCs to nucleate MTs under conditions where soluble tubulin concentration is elevated due to MT disassembly. The efficiency of MT nucleation, measured by staining for the MT plus-end tracking protein EB1, followed the same trend as centrosomal MT density, although the effects of the various knockouts were more pronounced (Fig. 1d,e; in 7KO and 8KO cells with separated centrosomes, the values for the two centrosomal asters were summed). We conclude that multiple PCM proteins redundantly regulate formation of a part of centrosomal MT population.

### PCM components control distinct centrosomal γ-TuRC pools

Since NIN and AKNA knockouts in 7KO and 8KO cells had little effect on MT density compared to 6KO cells, we first focused our subsequent analyses on 6KO cells, because they went through fewer rounds of genetic modification and selection. The intensities of PCM proteins NEDD1 and CEP192, as well as the centriole wall marker CEP295 ^68–70^ were reduced in 6KO cells compared to knockouts of AKAP450/CAMSAP2/CDK5RAP2/MMG or p53/PCNT (Fig. S1f,g). Ultrastructure Expansion Microscopy (U-ExM) ^71^ demonstrated that in 6KO cells, centrioles were significantly longer, sometimes had irregular shape and appeared thinner (Fig. 1g,h). Such defects were not observed in the p53/PCNT 2KO cells (Fig. 1g,h), suggesting that these abnormalities are caused by loss of multiple PCM components. Prolonged mitosis in 6KO cells (Fig. S1h,i) possibly contributed to the abnormal centriole elongation, as described previously ^72^. CEP192 tightly decorated the entire length of the centriole wall, whereas CEP295 was confined to its proximal region, as described previously ^46, 47, 69, 73, 74^ (Fig. 1g,h). The distribution of CEP192 and CEP295 was unchanged in p53/PCNT 2KO or 6KO cells (Fig. 1g,h), suggesting that the reduced signal of these proteins in 6KO cells is likely due to defects in centriole morphology rather than a direct dependence on the PCM proteins that were knocked out.

Centrosomal γ-tubulin accumulation was diminished in all studied knockouts, and the extent of reduction correlated with the efficiency of MT nucleation, with one notable exception: the knockout of AKNA in 8KO cells led to a significant loss of γ-tubulin intensity compared to 6KO and 7KO cells, whereas centrosomal MT nucleation and density were very similar in these knockouts (Fig. 1a-f). U-ExM analysis showed that the γ-tubulin pool in centriole lumen was not affected in 6KO, 7KO or 8KO cells (Fig. 1i). In contrast, pericentriolar γ-tubulin signal was strongly reduced in these cells: specifically, the cloud-like signal associated with the proximal centriole end was largely lost, while the signal enveloping the centriole wall was still present (Fig. 1i-k). We also observed γ-tubulin in a nine-fold symmetric pattern at subdistal appendages of the mother centriole (Fig. 1i). These signals were preserved in both 6KO and 7KO cells, but abolished in 8KO cells (Fig. 1i), indicating that AKNA, but not NIN, is responsible for recruiting γ-TuRC to subdistal appendages.

Together, these data demonstrate that cells where PCNT, AKAP450, CDK5RAP2 and MMG are knocked out lack γ-tubulin accumulation in the PCM around the proximal centriole end. This γ-TuRC pool contributes to MT nucleation after nocodazole washout. In the absence of these PCM proteins, NIN and AKNA do not support MT formation, although AKNA tethers γ-TuRC to subdistal appendages of the mother centriole independently of the abovementioned PCM components.

### NEDD1 and ch-TOG control centrosomal MT organization independently of PCNT, AKAP450, CDK5RAP2 and MMG

To study the origin of remaining MTs, we focused on the contribution of CEP192, NEDD1 and ch-TOG, which cooperate in nucleating MTs in mitotic *Xenopus* extracts and participate in centrosomal MT nucleation in interphase cells ^21, 43–46, 52^. Since these three proteins are essential for mitosis and cannot be stably knocked out, we used siRNA-mediated depletion. To achieve optimal reduction in protein levels (Fig. 2a,b), we followed a protocol whereby cells transfected with siRNAs twice (on day 1 and day 4) were arrested in interphase by thymidine block, and processed for immunofluorescence staining on day 7 after the first transfection.

**Figure 2.**
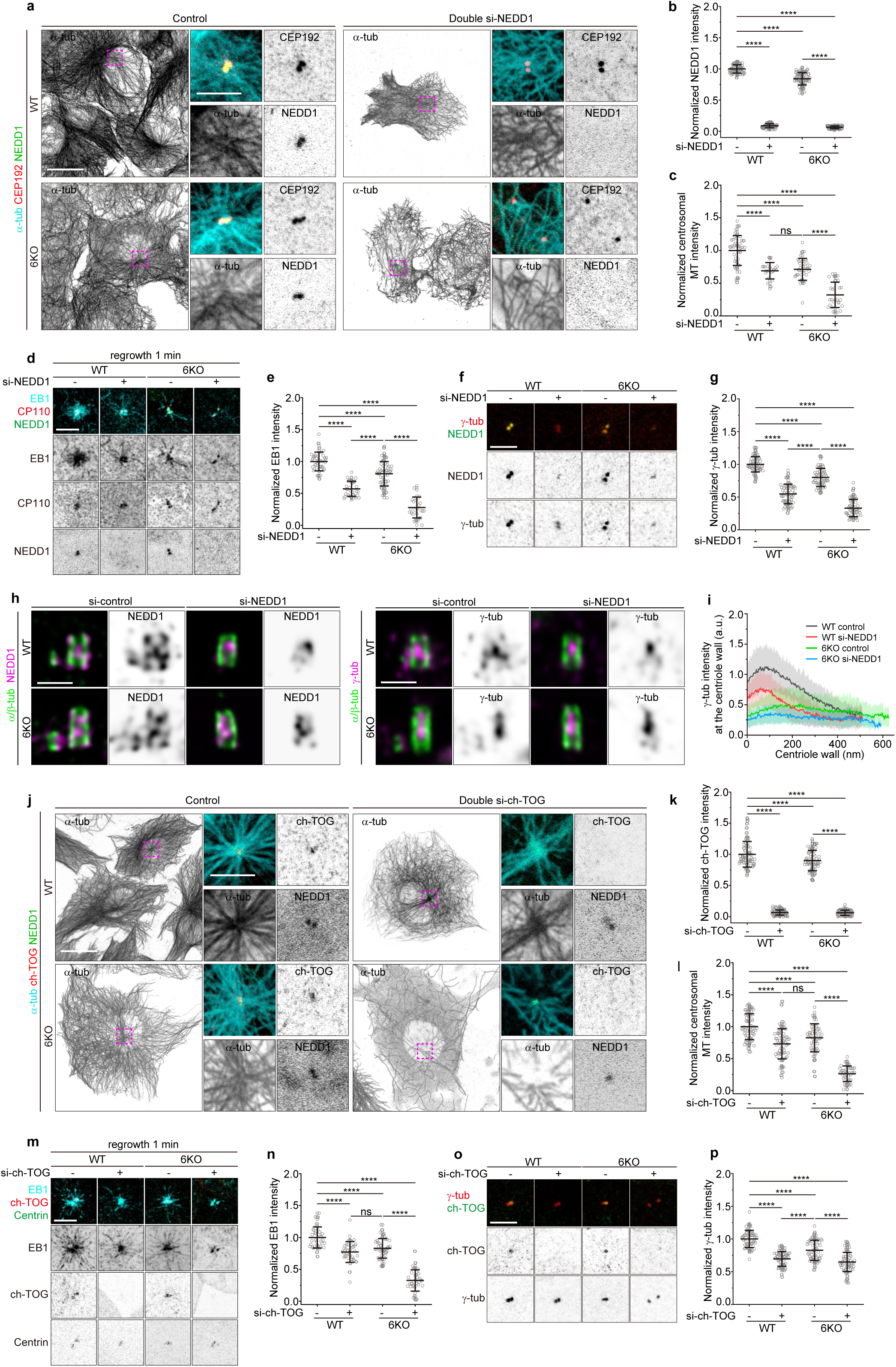
NEDD1 and ch-TOG control centrosomal MT organization independently of PCNT, AKAP450, CDK5RAP2 and MMG. **a,** Representative immunofluorescence images of RPE1 wild type and 6KO cells double-depleted of NEDD1, stained for α-tubulin (MT, cyan), CEP192 (red) and NEDD1 (green). Enlargements show merged and single channels of the boxed regions. Scale bar: 20 µm (main image); 5 µm (enlargement). **b,c,** Quantification of NEDD1 intensity per centrosome (mean ± s.d., **b**) and centrosomal MT intensity per cell (mean ± s.d., **c**) from the experiments shown in **a**. Values are normalized to the wild type average. The numbers of cells analyzed (n) for each condition from three independent experiments are: for NEDD1 (**b**): WT control (n = 78), WT si-NEDD1 (n = 78), 6KO control (n = 78), 6KO si-NEDD1 (n = 78); for centrosomal MT (**c**): WT control (n = 49), WT si-NEDD1 (n = 23), 6KO control (n = 37), 6KO si-NEDD1 (n = 28). **d,** Representative immunofluorescence images of RPE1 wild type and 6KO cells double-depleted of NEDD1, showing MT regrowth 1 min after nocodazole washout. Cells were stained for EB1 (cyan), CP110 (red) and NEDD1 (green). Scale bar: 5 µm. **e,** Quantification of centrosomal EB1 intensity per cell (mean ± s.d.) from the experiments shown in **d**. Values are normalized to the wild type average. The numbers of cells analyzed (n) for each condition from three independent experiments are: WT control (n = 44), WT si-NEDD1 (n = 34), 6KO control (n = 57), 6KO si-NEDD1 (n = 27). **f,** Representative immunofluorescence images of RPE1 wild type and 6KO cells double-depleted of NEDD1, stained for γ-tubulin (red) and NEDD1 (green). Scale bar: 5 µm. **g,** Quantification of γ-tubulin intensity per centrosome (mean ± s.d.) from the experiments shown in **f**. Values are normalized to the wild type average. The numbers of cells analyzed (n) for each condition from three independent experiments are: WT control (n = 59), WT si-NEDD1 (n = 59), 6KO control (n = 59), 6KO si-NEDD1 (n = 59). **h,** Representative U-ExM images of centrioles from RPE1 wild type and 6KO cells double-depleted of NEDD1, stained for α- and β-tubulin (green), NEDD1 (magenta, left) and γ-tubulin (magenta, right). Scale bar: 0.5 µm. **i,** Quantification of γ-tubulin intensity (mean ± s.d.) along the centriole from the proximal to the distal end in RPE1 wild type and 6KO cells double-depleted of NEDD1 (shown in **h**). The numbers of centrioles analyzed (n) for each condition from three independent experiments are: WT control, black, n = 38; WT si-NEDD1, red, n = 38; 6KO control, green, n = 43; 6KO si-NEDD1, blue, n = 24. **j,** Representative immunofluorescence images of RPE1 wild type and 6KO cells double-depleted of ch-TOG and stained for α-tubulin (MT, cyan), ch-TOG (red) and NEDD1 (green). Enlargements show merged and single channels of the boxed regions. Scale bar: 20 µm (main image); 5 µm (enlargement). **k, l,** Quantification of ch-TOG intensity per centrosome (mean ± s.d., **k**) and centrosomal MT intensity per cell (mean ± s.d., **l**) from the experiments shown in **j**. Values are normalized to the wild type average. The numbers of cells analyzed (n) for each condition from three independent experiments are: for ch-TOG (**k**): WT control (n = 76), WT si-ch-TOG (n = 76), 6KO control (n = 75), 6KO si-ch-TOG (n = 76); for centrosomal MT (**l**): WT control (n = 67), WT si-ch-TOG (n = 68), 6KO control (n = 53), 6KO si-ch-TOG (n = 45). **m,** Representative immunofluorescence images of RPE1 wild type and 6KO cells double-depleted of ch-TOG, showing MT regrowth 1 min after nocodazole washout. Cells were stained for EB1 (cyan), ch-TOG (red) and centrin (green). Scale bar: 5 µm. **n,** Quantification of centrosomal EB1 intensity per cell (mean ± s.d.) from the experiments shown in **m**. Values are normalized to the wild type average. The numbers of cells analyzed (n) for each condition from three independent experiments are: WT control (n = 36), WT si-ch-TOG (n = 39), 6KO control (n = 49), 6KO si-ch-TOG (n = 34). **o,** Representative immunofluorescence images of RPE1 wild type and 6KO cells double-depleted of ch-TOG, stained for γ-tubulin (red) and ch-TOG (green). Scale bar: 5 µm. **p,** Quantification of γ-tubulin intensity per centrosome (mean ± s.d.) from the experiments shown in **o**. Values are normalized to the wild type average. The numbers of cells analyzed (n) for each condition from three independent experiments are: WT control (n = 59), WT si-ch-TOG (n = 59), 6KO control (n = 59), 6KO si-ch-TOG (n = 59). Statistical significance was determined using one-way ANOVA with Tukey’s post hoc test for multiple comparisons (****P < 0.0001; ns, not significant, P > 0.05).

Depletion of NEDD1 abolished procentriole formation, as described previously ^75^, but had no effect on CEP295 or CEP192 (Fig. S2a-e). This indicates that NEDD1 does not contribute to centrosomal recruitment of CEP192. However, NEDD1 depletion in wild type cells reduced centrosomal MT density by 29%, comparable to the reduction observed in 6KO cells (Fig. 2c). When NEDD1 was depleted in 6KO cells, we observed an average reduction of 68% compared to wild type cells, with ~90% MTs lost in some cells (Fig. 2c). Analysis of MT regrowth after nocodazole washout followed the same pattern, although in this assay, the effect of NEDD1 depletion was stronger than the effect of PCM protein loss in 6KO cells (Fig. 2d,e; note that nocodazole washout data shown in each figure were collected simultaneously and can be compared with each other, but vary between figures, because even subtle differences in temperature and washout time affect EB1 signal). Centrosomal accumulation of γ-TuRC was also strongly affected by NEDD1 depletion (Fig. 2f,g); however, the reduction of γ-tubulin signal in 6KO cells depleted of NEDD1 was somewhat less profound than the reduction in MT density. U-ExM-based imaging explained this observation: despite sequential depletion, the intraluminal NEDD1 and γ-tubulin signal was still present in most cells, contrary to what was previously observed in U2OS cells ^46^. Quantification showed that NEDD1 depletion reduced intraluminal NEDD1 and γ-tubulin levels, although the reductions were modest (Fig. 2h, Fig. S2f,g). In wild type cells, NEDD1 loss strongly reduced the γ-tubulin pool present at the centriole wall, but not in the PCM surrounding the proximal centriole end (Fig. 2h,i). These data suggest that NEDD1 controls interphase γ-TuRC recruitment independently of PCNT, AKAP450, CDK5RAP2 and MMG.

Quantification of the centrosomal ch-TOG signal showed that it was mildly reduced (by ~10%) in 6KO cells compared to wild type cells, whereas NEDD1 depletion diminished the signal of ch-TOG almost by half in both wild type and 6KO cells (Fig. S2h,i). Similar to NEDD1 depletion, ch-TOG depletion did not affect the signals of CEP192 and CEP295 (Fig. S2k-m), whereas centrosomal NEDD1 levels in ch-TOG-depleted cells were decreased by ~25% in both wild type and 6KO cells (Fig. 2j, Fig. S2j). The depletion of ch-TOG reduced centrosomal MT density, nucleation and γ-tubulin recruitment in a pattern similar to NEDD1 depletion (Fig. 2j-p). These data indicate that NEDD1 and ch-TOG are interdependent for centrosomal recruitment and suggest that the two proteins cooperate in γ-TuRC recruitment and MT nucleation.

### CEP192 promotes centriole maintenance and MT nucleation in the absence of centrioles

We next used siRNA-mediated depletion to address the function of CEP192. A sequential double CEP192 depletion was lethal, and therefore, we used a single round of siRNA transfection. MT density around the centrosome and centrosomal MT nucleation were significantly diminished in CEP192-depleted cells, with the overall reduction much stronger in 6KO than in wild type cells (Fig. 3a-e). U-ExM confirmed that CEP192 specifically localizes to centriole wall (Fig. 3f,g). CEP192 knockdown reduced NEDD1 signal to a similar degree, by ~60%, in wild type and 6KO cells (Fig. 3h,i), and U-ExM confirmed that this reduction corresponds to the NEDD1 signal at the centriole wall, whereas NEDD1 enrichment within the PCM around proximal centriole end was minor (Fig. 3j,k). ch-TOG signal was also reduced after CEP192 depletion, and the same was true for γ-tubulin (Fig. 3a,h,l,m). U-ExM staining showed that the effect of CEP192 knockdown on γ-tubulin distribution was qualitatively similar to that of NEDD1 – it affected the γ-tubulin pool present along the centriole wall but not in the PCM around the centriole (Fig. 3n,o).

**Figure 3.**
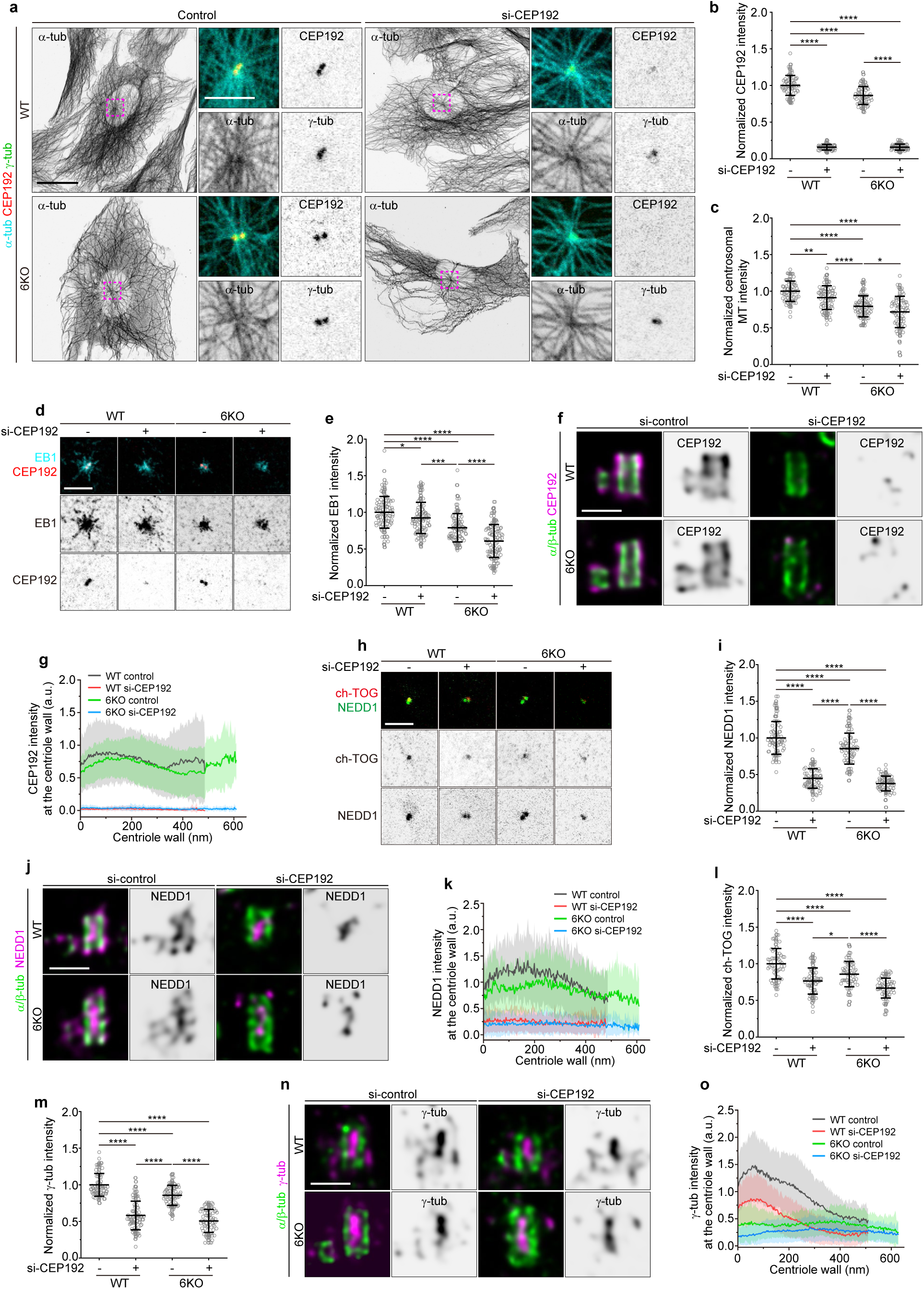
CEP192 controls centrosomal MT abundance by recruiting NEDD1 and ch-TOG. **a,** Representative immunofluorescence images of RPE1 wild type and 6KO cells depleted of CEP192, stained for α-tubulin (MT, cyan), CEP192 (red) and γ-tubulin (green). Enlargements show merged and single channels of the boxed regions. Scale bar: 20 µm (main image); 5 µm (enlargement). **b, c,** Quantification of CEP192 intensity per centrosome (mean ± s.d., **b**) and centrosomal MT intensity per cell (mean ± s.d., **c**) from the experiments shown in **a**. Values are normalized to the wild type average. The numbers of cells analyzed (n) for each condition from three independent experiments are: for CEP192 (**b**): WT control (n = 69), WT si-CEP192 (n = 70), 6KO control (n = 70), 6KO si-CEP192 (n = 70); for centrosomal MT (**c**): WT control (n = 74), WT si-CEP192 (n = 75), 6KO control (n = 82), 6KO si-CEP192 (n = 75). Statistical significance was determined using one-way ANOVA with Tukey’s post hoc test for multiple comparisons (* *P* = 0.0198; \*\**P* = 0.0073; \*\*\*\**P* < 0.0001). **d,** Representative immunofluorescence images of RPE1 wild type and 6KO cells depleted of CEP192, showing MT regrowth 1 min after nocodazole washout. Cells were stained for EB1 (cyan) and CEP192 (red). Scale bar: 5 µm. **e,** Quantification of centrosomal EB1 intensity per cell (mean ± s.d.) from the experiments shown in **d**. Values are normalized to the wild type average. The numbers of cells analyzed (n) for each condition from three independent experiments are: WT control (n = 101), WT si-CEP192 (n = 100), 6KO control (n = 99), 6KO si-CEP192 (n = 103). Statistical significance was determined using one-way ANOVA with Tukey’s post hoc test for multiple comparisons (* *P* = 0.0135; \*\*\**P* = 0.0007; \*\*\*\**P* < 0.0001). **f,** Representative U-ExM images of centrioles from RPE1 wild type and 6KO cells depleted of CEP192, stained for α- and β-tubulin (green) and CEP192 (magenta). Scale bar: 0.5 µm. **g,** Quantification of CEP192 intensity (mean ± s.d.) along the centriole from proximal to distal end in RPE1 wild type and 6KO cells depleted of CEP192 (shown in **f**). The numbers of centrioles analyzed (n) for each condition from three independent experiments are: WT control, black, n = 27; WT si-CEP192, red, n = 24; 6KO control, green, n = 39; 6KO si-CEP192, blue, n =24. **h,** Representative immunofluorescence images of RPE1 wild type and 6KO cells depleted of CEP192, stained for ch-TOG (red) and NEDD1 (green). Scale bar: 5 µm. **i, l,** Quantification of NEDD1 intensity per centrosome (mean ± s.d., **i**) and ch-TOG intensity per centrosome (mean ± s.d., **l**) from the experiments shown in **h**. Values are normalized to the wild type average. The numbers of cells analyzed (n) for each condition from three independent experiments are: for NEDD1 (**i**): WT control (n = 71), WT si-CEP192 (n = 71), 6KO control (n = 71), 6KO si-CEP192 (n = 71); for ch-TOG (**l**): WT control (n = 60), WT si-CEP192 (n = 65), 6KO control (n = 60), 6KO si-CEP192 (n = 60). Statistical significance was determined using one-way ANOVA with Tukey’s post hoc test for multiple comparisons (* *P* = 0.0166; \*\*\*\**P* < 0.0001). **j,** Representative U-ExM images of centrioles from RPE1 wild type and 6KO cells depleted of CEP192, stained for α- and β-tubulin (green) and NEDD1 (magenta). Scale bar: 0.5 µm. **k,** Quantification of NEDD1 intensity (mean ± s.d.) along the centriole from proximal to distal end in RPE1 wild type and 6KO cells depleted of CEP192 (shown in **j**). The numbers of centrioles analyzed (n) for each condition from three experiments are: WT control, black, n = 34; WT si-CEP192, red, n = 52; 6KO control, green, n = 37; 6KO si-CEP192, blue, n = 58. **m,** Quantification of γ-tubulin intensity per centrosome (mean ± s.d.) from the experiments shown in **a**. Values are normalized to the wild type average. The numbers of cells analyzed (n) for each condition from three independent experiments are: WT control (n = 75), WT si-CEP192 (n = 76), 6KO control (n = 76), 6KO si-CEP192 (n = 76). Statistical significance was determined using one-way ANOVA with Tukey’s post hoc test for multiple comparisons (\*\*\*\**P* < 0.0001). **n,** Representative U-ExM images of centrioles from RPE1 wild type and 6KO cells depleted of CEP192, stained for α- and β-tubulin (green) and γ-tubulin (magenta). Scale bar: 0.5 µm. **o,** Quantification of γ-tubulin intensity (mean ± s.d.) along the centriole from proximal to distal end in RPE1 wild type and 6KO cells depleted of CEP192 (shown in **n**). The numbers of centrioles analyzed (n) for each condition from three independent experiments are: WT control, black, n = 43; WT si-CEP192, red, n = 48; 6KO control, green, n = 39; 6KO si-CEP192, blue, n = 50.

We noted that the effects of CEP192 were weaker than those of NEDD1 depletion, possibly because CEP192 knockdown was incomplete. To achieve a more complete CEP192 depletion, we turned to inducible knockout. We used a lentiviral vector, which carries hU6 promoter for expressing a single guide RNA (sgRNA) targeting the CEP192-encoding gene, and two additional protein expression cassettes allowing Doxycycline-inducible expression of Cas9 and cytosolic GFP, and puromycin selection of transduced cells ^76^ (Fig. S3a). Transduced cells were selected using puromycin; after ~3 days of puromycin selection, Cas9 expression was induced by doxycycline addition for four days. During the last day, cells were blocked in the S phase by thymidine addition, because we observed significant cell death after the induction of CEP192 knockout, likely due to defects in cell cycle progression. Since not all cells were transduced, we selected for analysis only the cells which expressed GFP and which showed very strong loss of CEP192 based on immunofluorescence staining. Surrounding cells from the same coverslips that displayed no GFP signal were used as controls, both in the wild type and in 6KO background. In such cells, we could not detect other centriole markers, such as centrin and polyglutamylated tubulin (GT335), and centrosomal MT aster was also absent, indicating loss of the centrosomal MTOC (Fig. 4a).

**Figure 4.**
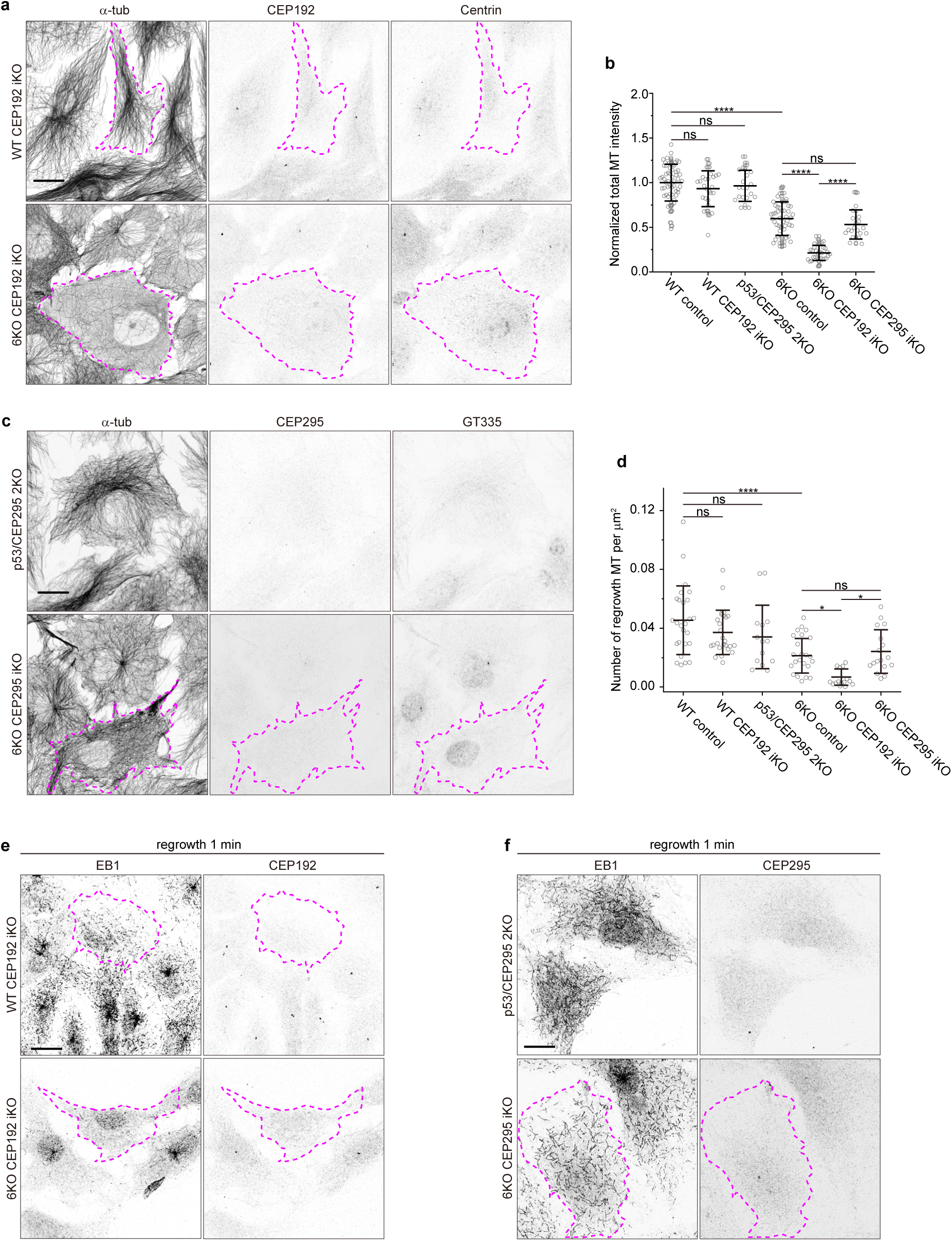
CEP192 is required for centriole maintenance and promotes MT nucleation in acentriolar cells. **a,** Representative immunofluorescence images of RPE1 wild type and 6KO cells with inducible knockout of CEP192, stained for α-tubulin (MT, grey), CEP192 (grey) and centrin (grey). CEP192 iKO cells are outlined with a dashed magenta line. Scale bar: 20 µm. **b,** Quantification of total MT intensity per cell (mean ± s.d.) from the experiments shown in **a** and **c**. Values are normalized to the wild type average. The numbers of cells analyzed (n) for each condition from three independent experiments are: WT control (n = 58), WT CEP192 iKO (n = 28), p53/CEP295 2KO (n = 21), 6KO control (n = 52), 6KO CEP192 iKO (n = 31), 6KO CEP295 iKO (n = 22). Statistical significance was determined using one-way ANOVA with Tukey’s post hoc test for multiple comparisons (\*\*\*\**P* < 0.0001; ns, not significant, *P* > 0.05). **c,** Representative immunofluorescence images of p53/CEP295 2KO RPE1 cells and 6KO cells with inducible knockout of CEP295, stained for α-tubulin (MT, grey), CEP295 (grey) and polyglutamylation (GT335, grey). CEP295 iKO cells are outlined with a dashed magenta line. Scale bar: 20 µm. **d,** Quantification of MT regrowth (number of EB1-positive MTs, mean ± s.d.) per square micrometer from the experiments shown in **e** and **f**. The numbers of cells analyzed (n) for each condition from three independent experiments are: WT control (n = 26), WT CEP192 iKO (n = 26), p53/CEP295 2KO (n = 14), 6KO control (n = 23), 6KO CEP192 iKO (n = 15), 6KO CEP295 iKO (n = 17). Statistical significance was determined using one-way ANOVA with Tukey’s post hoc test for multiple comparisons (*0.01< *P* < 0.05; \*\*\*\**P* < 0.0001; ns, not significant, *P* > 0.05). **e,** Representative immunofluorescence images of RPE1 wild type and 6KO cells with inducible knockout of CEP192, showing MT regrowth 1 min after nocodazole washout. Cells were stained for EB1 (grey) and CEP192 (grey). CEP192 iKO cells are outlined with a dashed magenta line. Scale bar: 20 µm. **f,** Representative immunofluorescence images of p53/CEP295 2KO RPE1 cells and 6KO cells with inducible knockout of CEP295, showing MT regrowth 1 min after nocodazole washout. Cells were stained for EB1 (grey) and CEP295 (grey). CEP295 iKO cells are outlined with a dashed magenta line. Scale bar: 20 µm.

Since CEP192 is required for centriole duplication ^47, 48^, centriole loss could occur due to division of cells that failed to duplicate their centriole. However, live imaging of cells with inducible knockout of CEP192 (CEP192 iKO) for 24 h after doxycycline induction stained with a live MT marker SiR-tubulin showed that MT disorganization occurred in interphase cells that did not go through mitosis, and fixation and staining of imaged cells confirmed absence of centrioles (Fig. S3b-e). These data indicate that inducible CEP192 knockout perturbed centriole maintenance in RPE1 cells. In the wild type background, acentriolar CEP192 iKO cells did not display any major loss in the overall MT density, in agreement with previous observations that MTs in acentriolar RPE1 cells can be maintained by the Golgi apparatus ^17, 18^. In contrast, in 6KO cells, inducible CEP192 knockout and the accompanying centriole loss dramatically reduced MT abundance (Fig. 4a,b).

To check whether this was due to the ability of the centrosome to promote MT formation by concentrating different MT-nucleating factors, we removed centrosomes by knocking out the centriole wall protein CEP295, which controls centrosome maturation and centriole maintenance ^68–70, 77^. As described previously ^70^, CEP295 could be stably knocked out in the p53 knockout background (Fig. 4c, Table S1). Similar to CEP192 iKO, centrosome loss in p53/CEP295 2KO cells did not affect overall MT density (Fig. 4b,c). In 6KO cells, constitutive CEP295 knockout was lethal, and we again turned to the inducible knockout strategy. Elimination of centrioles caused by CEP295 loss led to MT disorganization due to the absence of centrosomal MTOC, but MT density was maintained (Fig. 4b,c). Nocodazole washout assays showed that MT nucleation analyzed in the whole cell was lower in 6KO cells than in wild type cells but was not significantly affected by CEP295 knockout. In contrast, CEP192 loss did have a very profound effect on the number of newly nucleated MTs in 6KO cells (Fig. 4d-f). Together, these data show that CEP192 can promote MT formation in acentriolar 6KO cells. Since centrioles are absent in these cells, this suggests that CEP192 can participate in γ-TuRC activation in the cytoplasm, as illustrated by the random distribution of newly nucleated MT plus ends throughout the cell. It is thus the molecular activity of ch-TOG and associated proteins, and not their enrichment at the centrosome, that drives MT formation in interphase cells.

### Condensates formed by overexpressed CEP192 nucleate MTs in interphase cells

To further test the ability of CEP192 to activate MT nucleation in centrosome-independent context, we overexpressed it in RPE1 cells. mCherry-tagged CEP192 formed globular condensates, which recruited endogenous NEDD1, ch-TOG and γ-tubulin (Fig. 5a,b). Some other centrosomal and centriolar proteins, including CEP152, CEP295 and CEP135, were also present in these condensates. In contrast, other PCM proteins, such as PCNT, CDK5RAP2, AKAP450, NIN and AKNA, important for γ-TuRC recruitment to the Golgi, to the PCM around the proximal centriole end and the subdistal appendages, did not accumulate in CEP192 condensates (Fig. 5a,b, Fig. S4a). This is in line with our previous work, which showed that in centriole-depleted cells lacking Golgi MTs, PCNT, CDK5RAP2 and NIN could form a compact acentriolar MTOC (caMTOC) by dynein-driven self-assembly, while CEP192, NEDD1 and CEP152 did not participate in this process ^65^. In fact, CEP192 condensates and caMTOC have only two overlapping components, γ-tubulin and the MT polymerase ch-TOG (Fig. 5b). These data support the idea that CEP192 and PCNT together with CDK5RAP2 can self-organize into independent and biochemically distinct pathways of γ-TuRC recruitment and activation.

**Figure 5.**
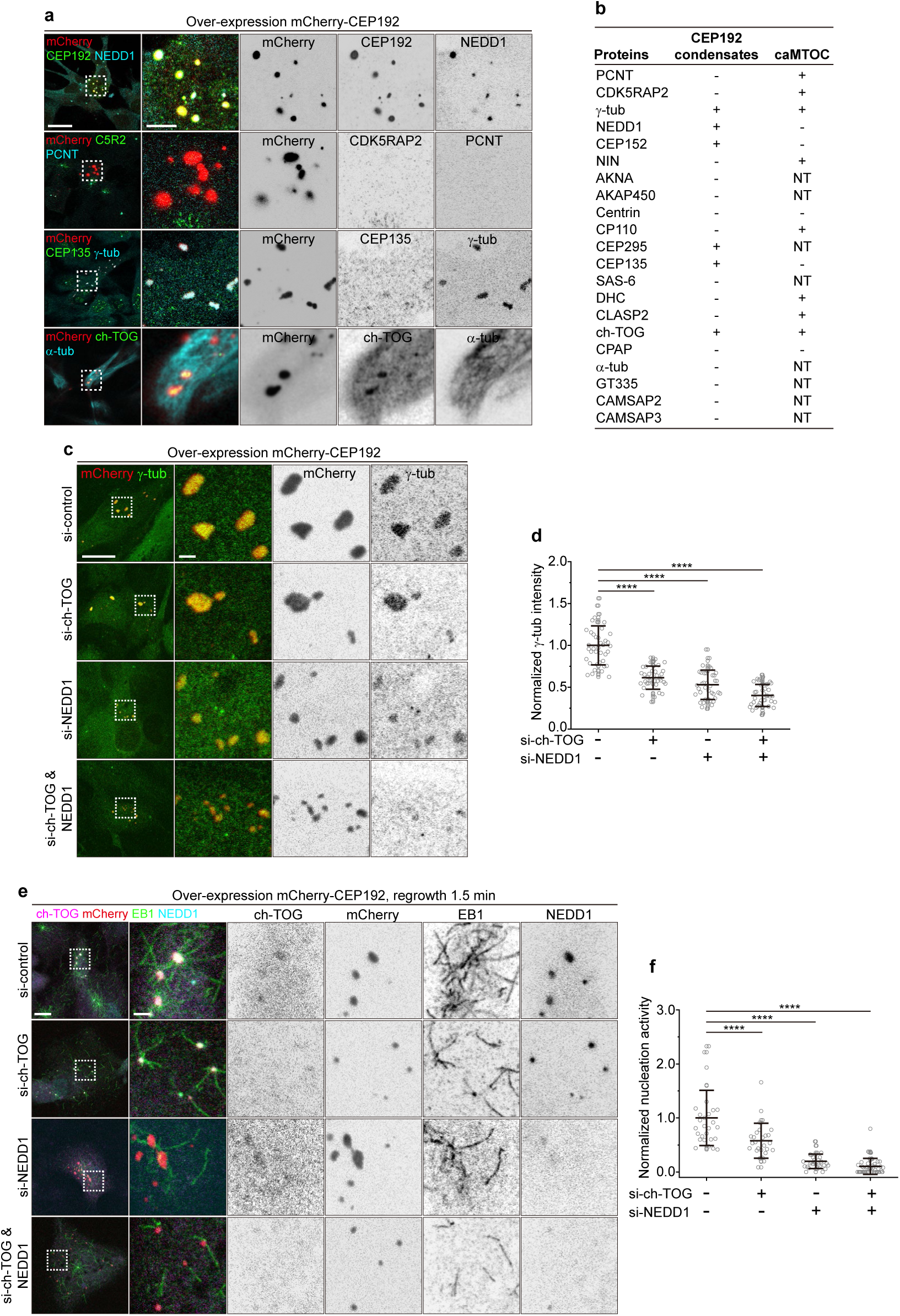
Condensates formed by overexpressed CEP192 nucleate MTs in interphase cells. **a,** Representative immunofluorescence images of mCherry-CEP192 condensates in transiently transfected wild type RPE1 cells stained for indicated proteins. Enlargements show merged and single channels of the boxed regions. Scale bar: 20 µm (main image); 5 µm (enlargement). **b,** Summary table of the indicated proteins present on mCherry-CEP192 condensates (**a** and **Fig. S4a**) and in the caMTOC in acentriolar cells ^65^. +, positive; −, negative; NT, not tested. **c,** Representative immunofluorescence images of mCherry-CEP192 condensates in RPE1 wild type cells. Cells were depleted of the indicated proteins and stained for mCherry (red) and γ-tubulin (green). Enlargements show merged and single channels of the boxed regions. Scale bar: 20 µm (main image); 5 µm (enlargement). **d,** Quantification of mean intensity of γ-tubulin (mean ± s.d.) in mCherry-CEP192 condensates from the experiments shown in **c**. Values are normalized to the control average. The numbers of condensates analyzed (n) for each condition from three independent experiments are: control (n = 40, 13 cells), si-ch-TOG (n = 40, 12 cells), si-NEDD1 (n = 40, 12 cells) and si-ch-TOG & si-NEDD1 (n = 40, 12 cells). Statistical significance was determined using one-way ANOVA with Tukey’s post hoc test for multiple comparisons (\*\*\*\**P* < 0.0001). **e,** Representative immunofluorescence images of mCherry-CEP192 condensates in RPE1 wild type cells, showing MT regrowth 1.5 min after nocodazole washout. Cells were depleted of indicated proteins and stained for EB1 (green), mCherry (red), ch-TOG (magenta) and NEDD1 (cyan). Enlargements show merged and single channels of the boxed regions. Scale bar: 20 µm (main image); 5 µm (enlargement). **f,** Nucleation activity (calculated as the number of EB1-positive MTs normalized to the mean intensity of γ-tubulin staining and CEP192 condensate area, mean ± s.d.) of mCherry-CEP192 condensates from the experiments shown in **e**. Values are normalized to the control average. The numbers of condensates analyzed (n) for each condition from three independent experiments are: control (n = 29, 15 cells), si-ch-TOG (n = 28, 20 cells), si-NEDD1 (n = 30, 16 cells) and si-ch-TOG & si-NEDD1 (n = 48, 17 cells). Statistical significance was determined using one-way ANOVA with Tukey’s post hoc test for multiple comparisons (\*\*\*\**P* < 0.0001).

To determine whether γ-tubulin recruitment to CEP192 condensates depends on NEDD1 and ch-TOG, we depleted these factors individually or in combination. NEDD1 depletion reduced ch-TOG accumulation in CEP192 condensates, whereas ch-TOG depletion had no effect on NEDD1 localization to these structures (Fig. S4b-f). However, both proteins were essential for the enrichment of γ-tubulin within the condensates (Fig. 5c,d). Nocodazole washout assays combined with protein depletion showed that CEP192 condensates can nucleate MTs in a manner dependent on NEDD1 and ch-TOG (Fig. 5e,f). These data suggest that CEP192 forms a scaffold that recruits NEDD1, and together, they recruit γ-TuRC and ch-TOG which nucleate MTs.

### In vitro reconstitution of MT nucleation by the complex of CEP192, NEDD1, ch-TOG and γ-TuRC

To clarify the interactions between CEP192, NEDD1, ch-TOG and γ-TuRC, we first performed pull-down assays from extracts of transfected HEK293T cells. Previous work has shown that the C-terminal part of CEP192 consists of eight ASPM–SPD-2–Hydin (ASH) domains, which have an immunoglobulin-like β-sandwich fold ^78–80^ (Fig. 6a, Fig. S5a). This part of the protein binds to CEP295 ^69^ and also drives formation of oligomeric PCM scaffold important for the mitotic function of CEP192 ^80^. The N-terminal part, which contains predicted unstructured regions interspersed with short helices, interacts with NEDD1 and γ-TuRC, Plk1, Plk4 and Aurora A kinases and potentially ch-TOG/XMAP215 ^47, 48, 52, 53, 55, 56, 81^. We mapped the NEDD1-binding domain of CEP192 to residues 768-930 (Fig. 6a, Fig. S5a,b). NEDD1 contains a C-terminal coiled-coil domain, which forms a tetrameric α-helical assembly that binds to γ-TuRC ^42^, and an N-terminal WD40 β-propeller domain, which targets NEDD1 to partners such as augmin ^82^ (Fig. 6a). Pull-down assays showed that similar to augmin, CEP192 interacts with the N-terminal part of NEDD1, leaving the C-terminus free for γ-TuRC recruitment (Fig. S5c-e). We also examined the interaction between CEP192 and ch-TOG and found that they interact through the N-terminal part of CEP192 (amino acids 1-768) and the C-terminal part of ch-TOG (Fig. 6a, Fig. S5f-h). This C-terminal region of ch-TOG is required for centrosomal targeting but not for binding to tubulin dimers ^21, 83^. Together, these data suggest that CEP192 can bring together ch-TOG and the NEDD1/γ-TuRC complex through non-overlapping binding sites.

**Figure 6.**
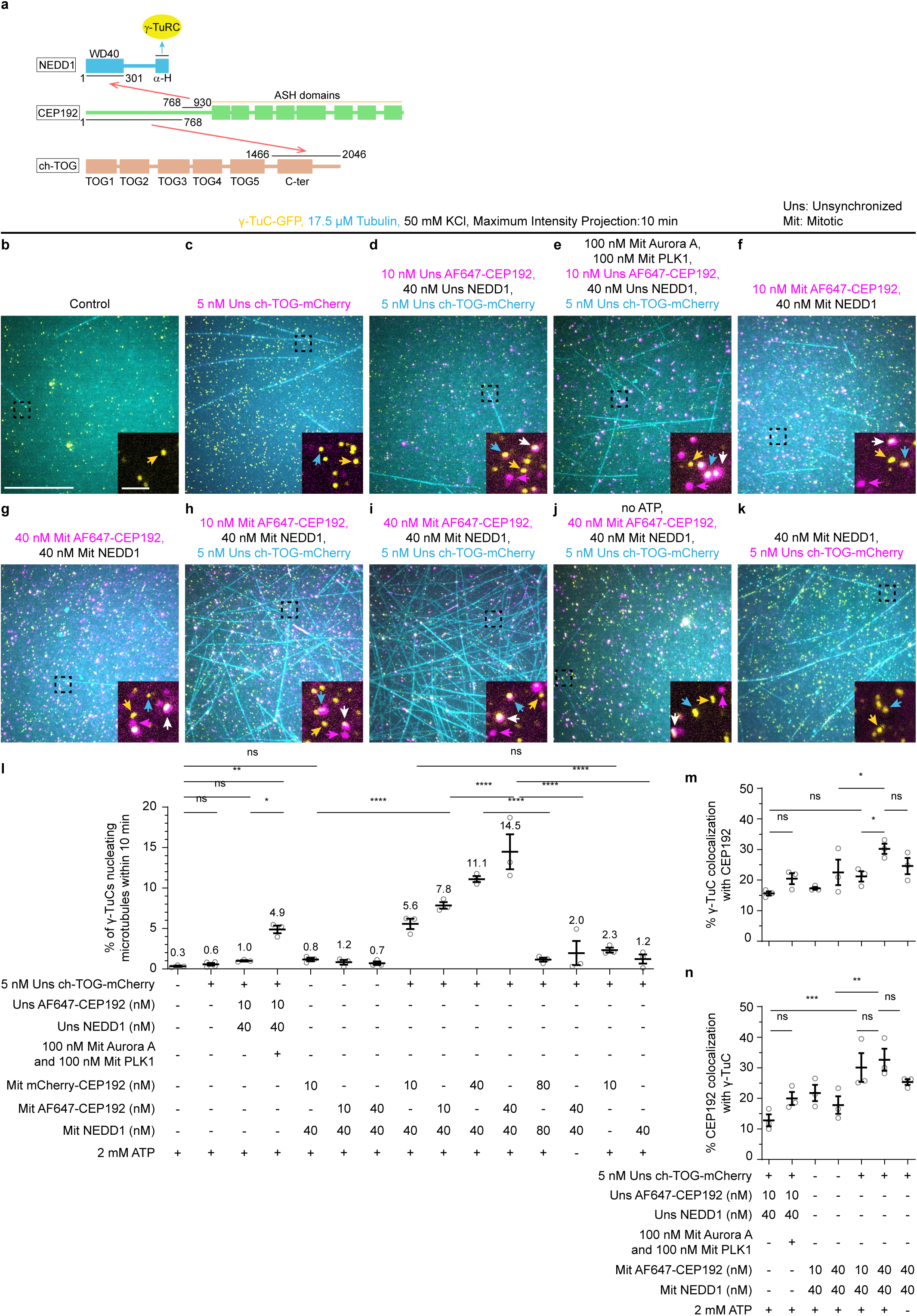
CEP192 and NEDD1 cooperate with ch-TOG to stimulate γ-TuC-mediated MT nucleation in vitro. **a,** Schematic representation of the domain structure of ch-TOG, NEDD1 and CEP192, and their interactions mapped by pull-down assays. **b-k,** Maximum intensity projections of 10 min videos, acquired after 10 min of incubation, showing MTs (cyan) nucleated from γ-TuCs (yellow, GCP3-GFP) under the indicated conditions: 17.5 μM tubulin control (**b**); 5 nM Uns ch-TOG-mCherry (magenta) alone (**c**); 10 nM Uns SNAP-AF647-CEP192 (magenta), 40 nM Uns NEDD1 and 5 nM Uns ch-TOG-mCherry (cyan) in the absence (**d**) or the presence (**e**) of 100 nM Mit Flag-Aurora A and 100 nM Mit SNAP-PLK1; 10 nM (**f**) or 40 nM (**g**) Mit SNAP-AF647-CEP192 (magenta) together with 40 nM Mit NEDD1; 10 nM (**h**) or 40 nM (**i**) Mit SNAP-AF647-CEP192 (magenta) together with 40 nM Mit NEDD1 and 5 nM Uns ch-TOG-mCherry (cyan); 40 nM Mit SNAP-AF647-CEP192 (magenta), 40 nM Mit NEDD1 and 5 nM Uns ch-TOG-mCherry (cyan) without ATP (**j**); 40 nM Mit NEDD1 together with 5 nM Uns ch-TOG-mCherry (magenta) (**k**). Magnification is the same in **b-k**. Insets are the magnified view of the black squares showing colocalization of γ-TuC shown in yellow with CEP192 or ch-TOG (in the assays without CEP192) shown in magenta. Yellow arrowheads in the insets indicate non-colocalizing γ-TuCs, cyan arrowheads indicate active γ-TuCs, magenta arrowheads indicate non-colocalizing CEP192 or ch-TOG, and white arrowheads indicate colocalizing γ-TuCs with CEP192 or ch-TOG. Magnification is the same in the insets in **b-k**. Scale bar: 20 µm, scale bar inset, 2 µm. Uns, proteins prepared from unsynchronized cells; Mit, proteins prepared from mitotic cells. **l**, Quantification of MT nucleation efficiency of γ-TuC in the indicated conditions. n=3 for all the assay conditions except for the assay with 5 nM Uns ch-TOG-mCherry (magenta) alone as represented in **c**, where n=4. n is the number of independent experiments performed per assay condition. Representative images are shown in **b-k** and **Fig. S6d-h**. The plot presents mean ± s.e.m., and each data point represents a single field of view for the given time point per experiment. Statistical significance was determined using one-way ANOVA with Tukey’s post hoc test for multiple comparisons (\**P* = 0.0473; \*\**P* = 0.0099; \*\*\*\**P* < 0.0001; ns, not significant, *P* > 0.05). **m,n,** Quantification of percentage colocalization (mean ± s.e.m.) of γ-TuC with SNAP-AF647-CEP192 (**m**) and vice versa (**n**) under given experimental conditions, represented in **d-j**. Each data point represents a single field of view for the given time point per experiment from which the percentage of γ-TuCs colocalizing with CEP192 or vice versa were quantified. n=3 for all assay conditions, where n is the number of independent experiments performed per assay condition. One-way ANOVA with uncorrected Fisher’s LSD tests was used to compare the means with each other (*0.01 < *P* < 0.05; \*\**P* = 0.0029; \*\*\**P* = 0.0009; ns, not significant, *P* > 0.05). In all the assay conditions, 2 mM ATP was added in the premixed γ-TuC solution before immobilization, as well as in the nucleation reaction mix with or without proteins, unless indicated otherwise.

To test whether these interactions are direct and whether the complex of CEP192, NEDD1, ch-TOG and γ-TuRC is sufficient to promote MT nucleation, we used in vitro assays with purified proteins. γ-TuRC was purified from a homozygous knock-in HEK293T cell line in which the GCP3-encoding gene was modified by a C-terminal insertion of GFP and a Twin-Strep-tag (SII) ^23^. Since not all purified complexes represent complete rings ^23^, they will be termed γ-tubulin complexes (γ-TuCs) for consistency with our previous study. Purified γ-TuCs were immobilized on coverslips using a biotinylated anti-GFP nanobody and used to observe MT nucleation in the presence of Rhodamine or HiLyte647-labeled tubulin by Total Internal Reflection Fluorescence (TIRF) microscopy (Fig. 6b-k).

As we and others have described previously ^5, 23, 84^, MT nucleation activity of purified γ-TuCs was low: within the first 10 min of observation after a 10 min incubation with the reaction mix, only 0.3% of γ-TuCs nucleated a MT (Fig. 6b,l). To test whether CEP192, NEDD1, and ch-TOG can activate γ-TuCs, we purified mCherry- or SNAP-AF647-tagged CEP192, untagged NEDD1, and ch-TOG-mCherry from unsynchronized HEK293T cells using a Twin-Strep-tag purification protocol, including washes with a high-ionic-strength buffer to improve protein purity (Fig. S6a). Mass spectrometry analysis of CEP192 and NEDD1 revealed low-level reciprocal co-purification: 0.3% NEDD1 in CEP192 preparation and 0.2% CEP192 in NEDD1 preparation, based on the intensity-based absolute quantification (iBAQ) values (Fig. S6b,c, Table S2 and S3). Some other centrosomal and PCM components were also detected (Fig. S6b,c, Table S2 and S3), but these co-purified low-abundance centrosome-associated proteins were mostly below 0.5%, except for Aurora A and PLK1 in CEP192 preparations (4.7% Aurora A and 0.7% PLK1, Fig. S6b). ch-TOG was absent from CEP192 and NEDD1 samples, and neither CEP192 nor NEDD1 were found in our ch-TOG preparation ^23^.

Next, we investigated whether the complex of CEP192, NEDD1, ch-TOG and γ-TuRC is sufficient to promote MT nucleation in our reconstitution assays. Our previous work showed that 200 nM ch-TOG could increase the nucleation frequency to ~10% and, when pre-incubated with γ-TuC before immobilizing it on a coverslip, could form large clusters that specifically sequestered γ-TuCs ^23^. To avoid these effects, we used ch-TOG-mCherry at a low concentration (5 nM) and added it directly to γ-TuCs immobilized on glass. At this concentration, ch-TOG did not enhance nucleation efficiency of γ-TuCs (only 0.6% γ-TuCs were activated, Fig. 6c,l), allowing us to assess potential synergistic effects with the CEP192-NEDD1 complex.

Because pre-incubation of CEP192 and NEDD1 with γ-TuCs did not lead to sequestration of γ-TuCs into clusters, we premixed 10 nM SNAP-AF647-CEP192 and 40 nM NEDD1 with γ-TuCs prior to their immobilization to maximize any stimulatory effect on γ-TuC nucleation efficiency. Since previous work has shown that CEP192 can be phosphorylated by mitotic kinases Aurora A and Plk1 ^52^, and Aurora A and PLK1 were present in our CEP192 preparation, we also included 2 mM ATP in our reconstitutions. However, we did not see any significant stimulation of MT nucleation in these conditions, as only 1% of γ-TuCs nucleated a MT (Fig. 6d,l).

Next, we supplemented these assays with 100 nM purified Flag-Aurora A and 100 nM SNAP-PLK1 (both without a fluorescent label, Fig. S6a), which were premixed with other assay components before attaching γ-TuCs to coverslips. This time, we observed a significant increase of nucleation frequency (Fig. 6e,l). Encouraged by these results, we purified CEP192 and NEDD1 from HEK293T cells synchronized in mitosis by a kinesin-5 inhibitor, S-trityl-L-cysteine (STLC) (Fig. S6a). Mitotic SNAP-AF647-CEP192 and NEDD1 again displayed low-level reciprocal co-purification (0.02% NEDD1 in CEP192 preparation and 0.5% CEP192 in NEDD1 preparation), as confirmed by our mass spectrometry results (Fig. S6b,c, Table S4 and S5). Moreover, Aurora A and PLK1 were also present in CEP192 preparation, though slightly less abundantly (2.1% Aurora A and 0.6% PLK1, Fig. S6b) compared to CEP192 purified from unsynchronized cells. This combination of CEP192 and NEDD1 still did not significantly activate MT nucleation by itself, even when CEP192 concentration was increased by four times from 10 to 40 nM (Fig. 6f,g,l, Fig. S6d). However, a very significant (7-fold) activation was observed in the presence of 5 nM ch-TOG (Fig. 6h,l, Fig. S6e). This effect was even stronger and peaked when CEP192 concentration was increased from 10 to 40 nM (Fig. 6i,l, Fig. S6f). Doubling the concentrations of CEP192 and NEDD1 to 80 nM each caused a strong reduction in γ-TuC nucleation efficiency (Fig. 6l, Fig. S6g). mCherry-CEP192 and SNAP-AF647-CEP192 versions activated γ-TuCs to a similar extent (see Fig. 6h,i for SNAP-tagged and Fig. S6e,f for mCherry-tagged CEP192). The stimulatory effect on MT nucleation efficiency of γ-TuC was dependent on ATP, since omitting ATP from our reconstitutions abolished γ-TuC activation (Fig. 6j,l). Furthermore, γ-TuC activation was also dependent on the complex of CEP192-NEDD1, as ch-TOG was not sufficient to strongly activate γ-TuC in the presence of NEDD1 or CEP192 alone (Fig. 6k,l, Fig. S6h).

Even at these low concentrations, we could observe very significant (up to ~30%) reciprocal colocalization between γ-TuC and CEP192 (Fig. 6m,n). When CEP192 and NEDD1 were prepared from unsynchronized cells, the colocalization between CEP192 and γ-TuC was slightly enhanced by pre-incubation with mitotic kinases (Fig. 6d,e,m), and it was even higher when CEP192 and NEDD1 were isolated from mitotic cells (Fig. 6h,i,m). Moreover, the addition of ch-TOG, prepared from unsynchronized cells, increased colocalization between γ-TuCs and CEP192 (Fig. 6f-i,m), in agreement with the data in Fig. 2o,p and Fig. 5c,d, and the previously described role of this MT polymerase in centrosomal γ-TuRC recruitment ^21^. Taken together, our biochemical and reconstitution data indicate that CEP192 can form complexes with NEDD1, γ-TuRC and ch-TOG and stimulate MT nucleation. This process strongly depends on regulatory phosphorylation, which promotes the interaction between these proteins.

## Discussion

In this study, we systematically dissected the contributions of centrosomal proteins involved in γ-TuRC recruitment and activation to MT formation in interphase cells. We used RPE1 cells because the major components of the Golgi-associated MTOC have already been characterized in this cell line ^17, 18^, facilitating analysis of centrosomal contribution to MT formation. Most of the proteins studied here have previously been knocked out or depleted individually or in smaller combinations, and in most cases, there was no or only a moderate effect on the MT density, because the inhibition of one pathway, for example, centrosomal nucleation, caused redistribution of γ-TuRC and its activators to non-centrosomal sites ^17, 18, 23^. This led to the view that different nucleation activators control the architecture of the MT network, but their importance for MT abundance in cells remained unclear. Here, we found that in interphase RPE1 cells, simultaneous knockout of CAMSAP2, AKAP450, PCNT, CDK5RAP2, MMG and CEP192 caused profound reduction of the MT density. CAMSAP2 is a minus end stabilizer which does not nucleate MTs but rather binds free MT minus ends or displaces γ-TuRC from MTs ^23, 85^, whereas the rest are γ-TuRC activators or scaffolds. Therefore, we conclude that in the absence of CAMSAP2-mediated MT minus-end stabilization, activation of γ-TuRC becomes essential to maintain MT density.

Our data indicate that γ-TuRC-binding proteins operate in interphase RPE1 cells in several distinct pathways. A significant pool of γ-TuRC resides in the PCM surrounding the proximal centriole end. This pool, as well as Golgi MTs, are lost in cells lacking PCNT, CDK5RAP2, AKAP450 and MMG. The reduction in centrosomal MT density in cells lacking all these proteins and CAMSAP2 (6KO cells) was somewhat stronger than in any of the cell lines missing their subsets (e.g., PCNT alone). This indicates that these proteins cooperate in the formation of PCM, which recruits and activates γ-TuRC. However, centrioles and centrosomes still form and function in 6KO cells, even though mitosis takes significantly longer, as expected from previous work ^32, 59, 86^. Moreover, 6KO cells display centriole defects, suggesting that the PCM missing in these cells has an impact on centriole biogenesis. Centriole defects were not observed in cells lacking only PCNT, the key factor responsible for centrosome recruitment of CDK5RAP2, suggesting that they are caused by the collective loss of several PCM components, similar to what has been described in *Drosophila* ^87^.

In contrast to PCM, γ-TuRC localization at the subdistal appendages was not affected in 6KO cells. Previous work implicated NIN in γ-TuRC docking and MT minus end anchoring at subdistal appendages ^38, 39^, but it turned out that in RPE1 cells this γ-TuRC pool depends on AKNA, in agreement with the reported AKNA function in developing neuronal cells ^37^. However, in RPE1 cells, NIN and AKNA do not contribute much to the overall MT organization. Cells missing CAMSAP2, AKAP450, PCNT, CDK5RAP2, MMG, NIN and AKNA on the background of p53 knockout were viable and contained functional centrosomes, even though centrosome linker was lost due to the absence of NIN, as described previously ^66, 67^. Whether AKNA can independently activate γ-TuRC or requires cofactors such as ch-TOG, which can also accumulate at subdistal appendages ^21^, remains to be determined in a cell type in which AKNA-dependent MT organization makes a larger functional contribution.

The remaining MT density in 6KO cells strongly relied on the complex of CEP192, NEDD1 and ch-TOG. This complex localizes along the centriole wall through CEP192, in agreement with previous work ^46–49^. The loss of either NEDD1 or ch-TOG had no effect on CEP192 localization, indicating that the recently described electrostatic interaction of NEDD1 with polyglutamylated MTs ^88^ does not contribute to centriole recruitment of CEP192. In contrast, the localization of NEDD1 and ch-TOG was partly interdependent, supporting previous work ^21^. Centrosome recruitment of NEDD1 and ch-TOG required CEP192 but was only weakly affected by the loss of PCNT, AKAP450, CDK5RAP2 and MMG. Conversely, the latter proteins could still form a functional PCM to activate γ-TuRC in cells depleted of CEP192, NEDD1 and ch-TOG. Therefore, unlike mitotic cells, where PCNT, CDK5RAP2 and CEP192 cooperate in spindle pole formation ^49, 59^, the interphase CEP192/NEDD1/ch-TOG pathway appears to act independently of PCNT and CDK5RAP2. This view is supported by the observation that PCNT does not colocalize with CEP192 and NEDD1 in acentriolar interphase cells ^65^ or in interphase condensates.

In addition to recruiting NEDD1 and γ-TuRC and participating in initiation of centriole biogenesis ^46–48^, centriolar localization of CEP192 serves another important function – it helps to maintain centriole integrity, as centrioles appeared to disassemble when CEP192 was strongly depleted by an inducible knockout. This effect has not been observed previously ^43, 46–48, 51^, possibly because both siRNA-mediated depletion and inducible protein degradation reduce CEP192 pool less profoundly, or due to cell type-specific differences.

In contrast to PCNT or NIN ^89, 90^, CEP192 does not seem to interact with cytoplasmic dynein, and in the absence of centrioles, it does not accumulate in one spot but redistributes to the cytoplasm. Importantly, the ability of CEP192 to activate γ-TuRC is not dependent on being concentrated at the centrosome, as it also increases MT density in acentriolar CEP295 knockout cells lacking CAMSAP2, AKAP450, PCNT, CDK5RAP2 and MMG. CEP192 can also activate single γ-TuRCs in vitro in a complex with NEDD1 and ch-TOG. Although this activity is clearly detectable in interphase cells, we were unable to recapitulate it in vitro in the absence of mitotic kinases, perhaps because it is quite low – even large CEP192 condensates nucleated only a few interphase MTs. In agreement with previous work in *Xenopus* extracts ^52^, the binding of CEP192 to γ-TuRC in the presence of NEDD1 and ch-TOG is strongly potentiated by mitotic kinases, for which CEP192 serves as a scaffold. Whether some additional, interphase-specific factors potentiate γ-TuRC activation in the context of CEP192/NEDD1/ch-TOG complex, or whether this depends on the weak activities of Aurora A and PLK1 in interphase cells remains to be determined.

The relative contributions of centrosomal MT nucleation pathways vary between cell types, and the method of protein inactivation can affect the conclusions. For example, acute depletion of PCNT mildly increased the number of MTs nucleated from centrosomes in U2OS cells ^43^, but decreased centrosome activity in our stable knockout RPE1 cells. In spite of these differences, a systematic dissection of γ-TuRC recruitment and activation pathways in a single cell type is valuable, because it allows determining their biochemical interdependence. For example, our data show that although ch-TOG is a general MT polymerase that directly interacts with and activates γ-TuRC ^22–24^, it does not seem essential for nucleating PCNT/CDK5RAP2-dependent centrosomal MTs. Whether γ-TuRCs activated through different pathways acquire distinct structures and functional properties — for example, by differentially regulating MT minus-end stability and release through associated proteins or mechanical forces, thereby generating MTs with distinct lattice architecture and post-translational modifications, remains to be determined.

The most important conclusion of our study is that MT formation in interphase mammalian cells critically depends on γ-TuRC activation. This finding supports the notion that intrinsic MT nucleation activity of γ-TuRC in cells is poor. Furthermore, although various MAPs can drive γ-TuRC-independent MT nucleation in vitro ^2^, and γ-TuRC-independent nucleation has been demonstrated in cells ^27, 91^, our results indicate that, at least in some types of interphase mammalian cells, γ-TuRC-dependent nucleation and subsequent anchoring is not merely kinetically favored but also presents the only alternative to CAMSAP-mediated MT minus-end stabilization. One possible reason why cells rely on this seemingly elaborate mechanism to generate MTs is the need to tightly control MT architecture. Whereas γ-TuRC nucleates MTs with a defined 13-protofilament structure ^9, 10^, spontaneously nucleated MTs exhibit substantial variability in protofilament number ^92, 93^. The knockout cell lines lacking multiple γ-TuRC regulators described here provide a powerful experimental system to further dissect MT formation pathways and explore the functional consequences of their disruption.

## Acknowledgements

We thank S. Lens (UMCU, the Netherlands), G. Kops (Hubrecht Institute, the Netherlands), A. Holland (Johns Hopkins University School of Medicine, USA) and D. Trono (EPFL, Switzerland) for sharing materials. This work was supported by China Scholarship Council scholarship to F. C., the European Research Council Synergy grant PushingCell (project number 101071793) and the Netherlands Organization for Scientific Research Spinoza prize to A.A. The authors acknowledge support from the Dutch Research Council-funded Netherlands Proteomics Centre through the National Roadmap for Large-scale Research Infrastructures programme X-Omics (project 184.034.019) to K.S. for the mass spectrometry analysis.

## Author contributions

Y.S. designed and performed experiments with cells, analyzed data and wrote the paper; D.R. designed and performed protein purifications and in vitro reconstitution experiments, analyzed data and wrote the paper; L.M.S. assisted with the expansion microscopy experiments; M.F.M. S. and K.E.S. performed, analyzed and supervised mass spectrometry experiments; B.J.K. generated recombinant constructs, S.T.K. performed experiments, Q.J.K. analyzed data; F.C. initiated the project, generated and characterized several knockout cell lines, developed and provided phenotypic validation, mechanistic analysis and supervision, and wrote the paper; A.A. coordinated the project and wrote the paper.

## Supplemental Figures

**Figure S1.**
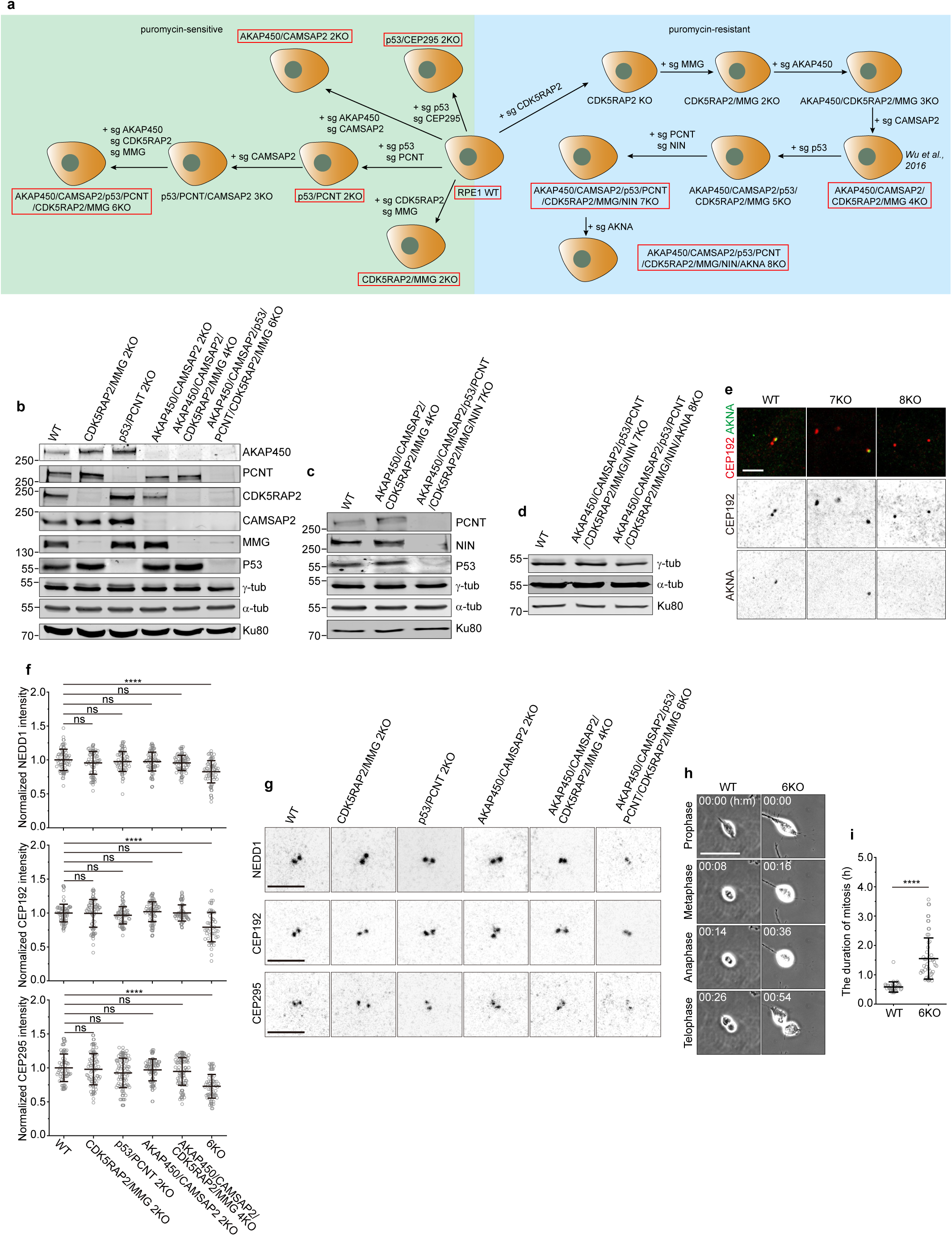
Characterization of RPE1 knockout cell lines. **a,** Schematic representation of the generation of RPE1 knockout cell lines. Cell lines used in this study are indicated by red boxes. **b-d,** Western blot of cell lysates from RPE1 wild type and indicated knockout cell lines, blotted using indicated antibodies. **e,** Representative immunofluorescence images of indicated RPE1 cell lines, stained for CEP192 (red) and AKNA (green). Scale bar: 5 µm. **f,** Quantification of NEDD1, CEP192 and CEP295 intensity per centrosome (mean ± s.d.) from the experiments shown in **g**. Values are normalized to the wild type average. The numbers of cells analyzed for each cell line (in the same order as the graph) are as follows: for NEDD1: 59, 54, 56, 63, 63 and 53; for CEP192: 64, 62, 62, 63, 62 and 43; for CEP295: 60, 67, 79, 81, 82 and 61. Each protein quantification was performed on data from three independent experiments. Statistical significance was determined using one-way ANOVA with Tukey’s post hoc test for multiple comparisons (\*\*\*\**P* < 0.0001; ns, not significant, *P* > 0.05). **g,** Representative immunofluorescence images of indicated RPE1 cell lines, stained for NEDD1 (grey, top), CEP192 (grey, middle) and CEP295 (grey, bottom). Scale bar: 5 µm. **h,** Representative phase-contrast images of RPE1 wild type and 6KO cell lines in mitosis. Scale bar: 50 µm. **i,** Quantification of mitotic duration (mean ± s.d.) in RPE1 wild type (n = 34) and 6KO (n = 40) cell lines from the experiments shown in **h**. n indicates the number of cells analyzed, collected from two independent experiments. Statistical significance was determined using one-way ANOVA with Tukey’s post hoc test for multiple comparisons (\*\*\*\**P* < 0.0001).

**Figure S2.**
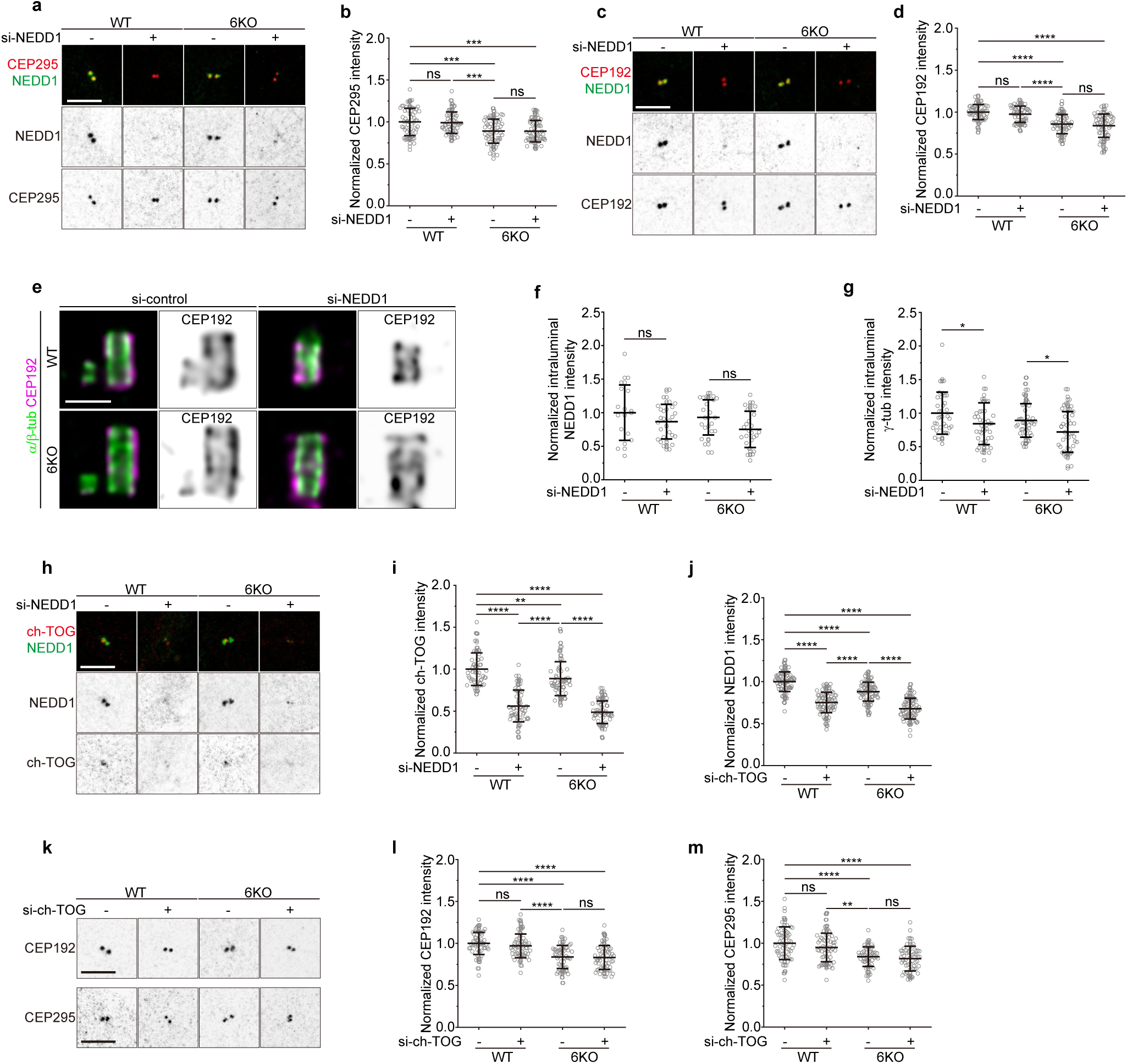
Depletion of NEDD1 and ch-TOG does not affect centrosomal levels of CEP192 and CEP295 at the centriole. **a,** Representative immunofluorescence images of RPE1 wild type and 6KO cells double-depleted of NEDD1, stained for CEP295 (red) and NEDD1 (green). Scale bar: 5 µm. **b,** Quantification of CEP295 intensity per centrosome (mean ± s.d.) from the experiments shown in **a**. Values are normalized to the wild type average. The numbers of cells analyzed (n) for each condition from three independent experiments are: WT control (n = 59), WT si-NEDD1 (n = 58), 6KO control (n = 59), 6KO si-NEDD1 (n = 59). Statistical significance was determined using one-way ANOVA with Tukey’s post hoc test for multiple comparisons (***0.0001 < *P* < 0.001; ns, not significant, *P* > 0.05). **c,** Representative immunofluorescence images of RPE1 wild type and 6KO cells double-depleted of NEDD1 and stained for CEP192 (red) and NEDD1 (green). Scale bar: 5 µm. **d,** Quantification of CEP192 intensity per centrosome (mean ± s.d.) from the experiments shown in **c**. Values are normalized to the wild type average. The numbers of cells analyzed (n) for each condition from three independent experiments are: WT control (n = 59), WT si-NEDD1 (n = 59), 6KO control (n = 59), 6KO si-NEDD1 (n = 59). Statistical significance was determined using one-way ANOVA with Tukey’s post hoc test for multiple comparisons (\*\*\*\**P* < 0.0001; ns, not significant, *P* > 0.05). **e,** Representative U-ExM images of centrioles from RPE1 wild type and 6KO cells double-depleted of NEDD1, stained for α- and β-tubulin (green) and CEP192 (magenta). Scale bar: 0.5 µm. **f,g,** Quantification of intraluminal NEDD1 (f) and γ-tubulin (g) intensity per centriole (mean ± s.d.) from the experiments shown in Fig. 2h. Values are normalized to the wild type average. The numbers of centrioles analyzed (n) for each condition from two (f) or three (g) independent experiments are: for NEDD1 (f): WT control (n = 17), WT si-NEDD1 (n = 29), 6KO control (n = 23), 6KO si-NEDD1 (n = 25); for γ-tubulin (g): WT control (n = 37), WT si-NEDD1 (n = 42), 6KO control (n = 45), 6KO si-NEDD1 (n = 43). Statistical significance was determined using one-way ANOVA with Tukey’s post hoc test for multiple comparisons (*0.01 < P < 0.05; ns, not significant, P > 0.05). **h,** Representative immunofluorescence images of RPE1 wild type and 6KO cells double-depleted of NEDD1, stained for ch-TOG (red) and NEDD1 (green). Scale bar: 5 µm. **i,** Quantification of ch-TOG intensity per centrosome (mean ± s.d.) from the experiments shown in **h**. Values are normalized to the wild type average. The numbers of cells analyzed (n) for each condition from three independent experiments are: WT control (n = 54), WT si-NEDD1 (n = 59), 6KO control (n = 56), 6KO si-NEDD1 (n = 59). Statistical significance was determined using one-way ANOVA with Tukey’s post hoc test for multiple comparisons (\*\**P* = 0.0066, \*\*\*\**P* < 0.0001). **j,** Quantification of NEDD1 intensity per centrosome (mean ± s.d.) from the experiments shown in Fig. 2j. Values are normalized to the wild type average. The numbers of cells analyzed (n) for each condition from three independent experiments are: WT control (n = 80), WT si-ch-TOG (n = 76), 6KO control (n = 78), 6KO si-ch-TOG (n = 79). Statistical significance was determined using one-way ANOVA with Tukey’s post hoc test for multiple comparisons (\*\*\*\**P* < 0.0001). **k,** Representative immunofluorescence images of RPE1 wild type and 6KO cells double-depleted of ch-TOG, stained for CEP192 (grey) and CEP295 (grey). Scale bar: 5 µm. **l,m,** Quantification of CEP192 intensity per centrosome (mean ± s.d., **l**) and CEP295 intensity per centrosome (mean ± s.d., **m**) from the experiments shown in **k**. Values are normalized to the wild type average. The numbers of cells analyzed (n) for each condition from three independent experiments are: for CEP192 (**l**): WT control (n = 58), WT si-ch-TOG (n = 58), 6KO control (n = 54), 6KO si-ch-TOG (n = 57); for CEP295 (**m**): WT control (n = 58), WT si-ch-TOG (n = 58), 6KO control (n = 55), 6KO si-ch-TOG (n = 58). Statistical significance was determined using one-way ANOVA with Tukey’s post hoc test for multiple comparisons (\*\**P* = 0.0019; \*\*\*\**P* < 0.0001; ns, not significant, *P* > 0.05).

**Figure S3.**
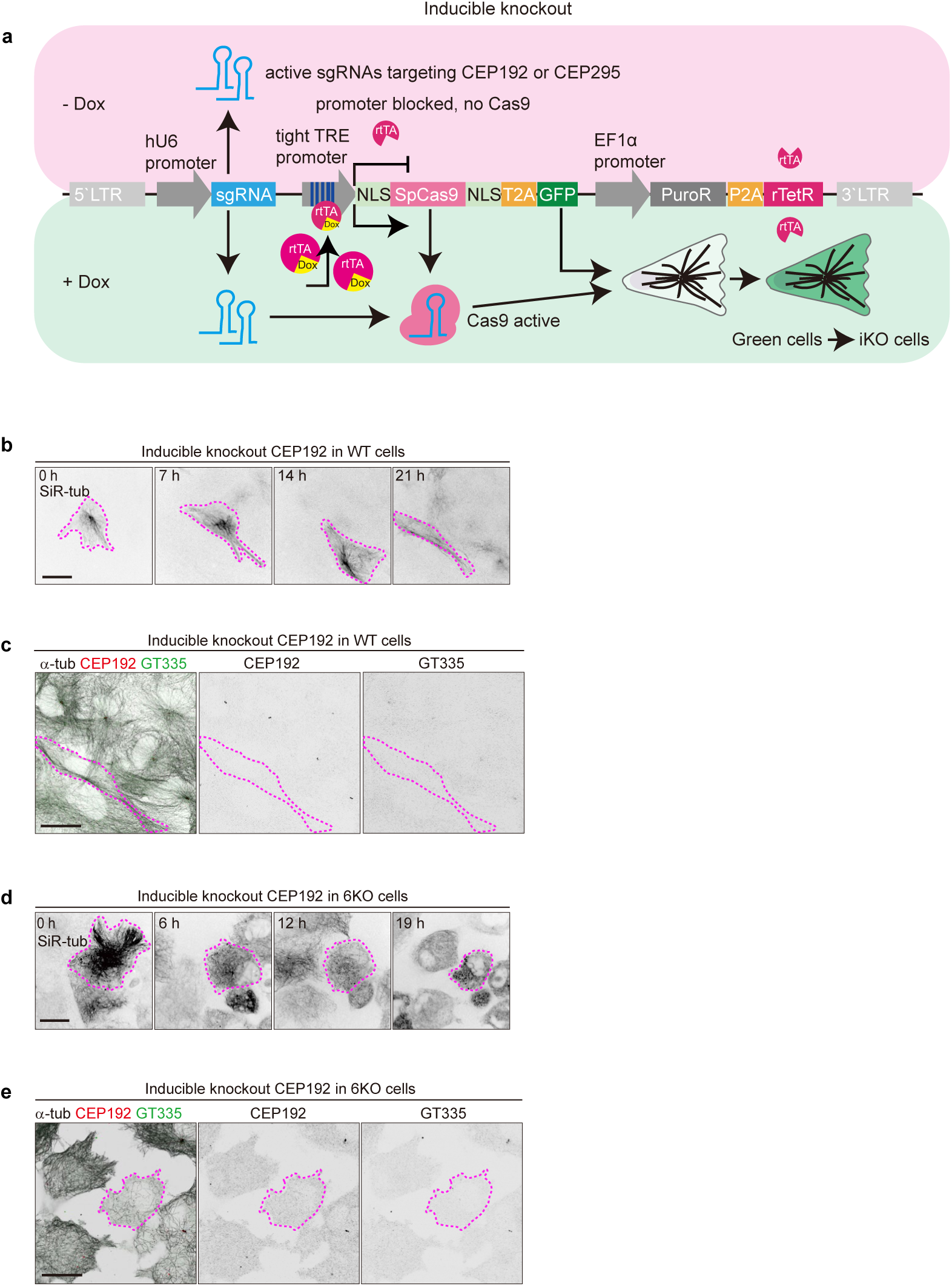
Characterization of CEP192 iKO cells. **a,** Scheme of the doxycycline-inducible knockout strategy. **b,** Time-lapse images of RPE1 wild type cells with inducible knockout of CEP192, labeled with 100 nM SiR-tubulin (grey). Scale bar: 20 µm. **c,** Immunofluorescence images of fixed cells shown in **b.** Cells were stained for α-tubulin (MT, grey), CEP192 (red) and polyglutamylation (GT335, green). CEP192 knockout cells are outlined with a dashed magenta line. Scale bar: 20 µm. **d,** Time-lapse images of RPE1 6KO cells with inducible knockout of CEP192, labeled with 100 nM SiR-tubulin (grey). Scale bar: 20 µm. **e,** Immunofluorescence images of fixed cells shown in **d**, Cells were stained for α-tubulin (MT, grey), CEP192 (red) and GT335 (green). CEP192 knockout cells are outlined with a dashed magenta line. Scale bar: 20 µm.

**Figure S4.**
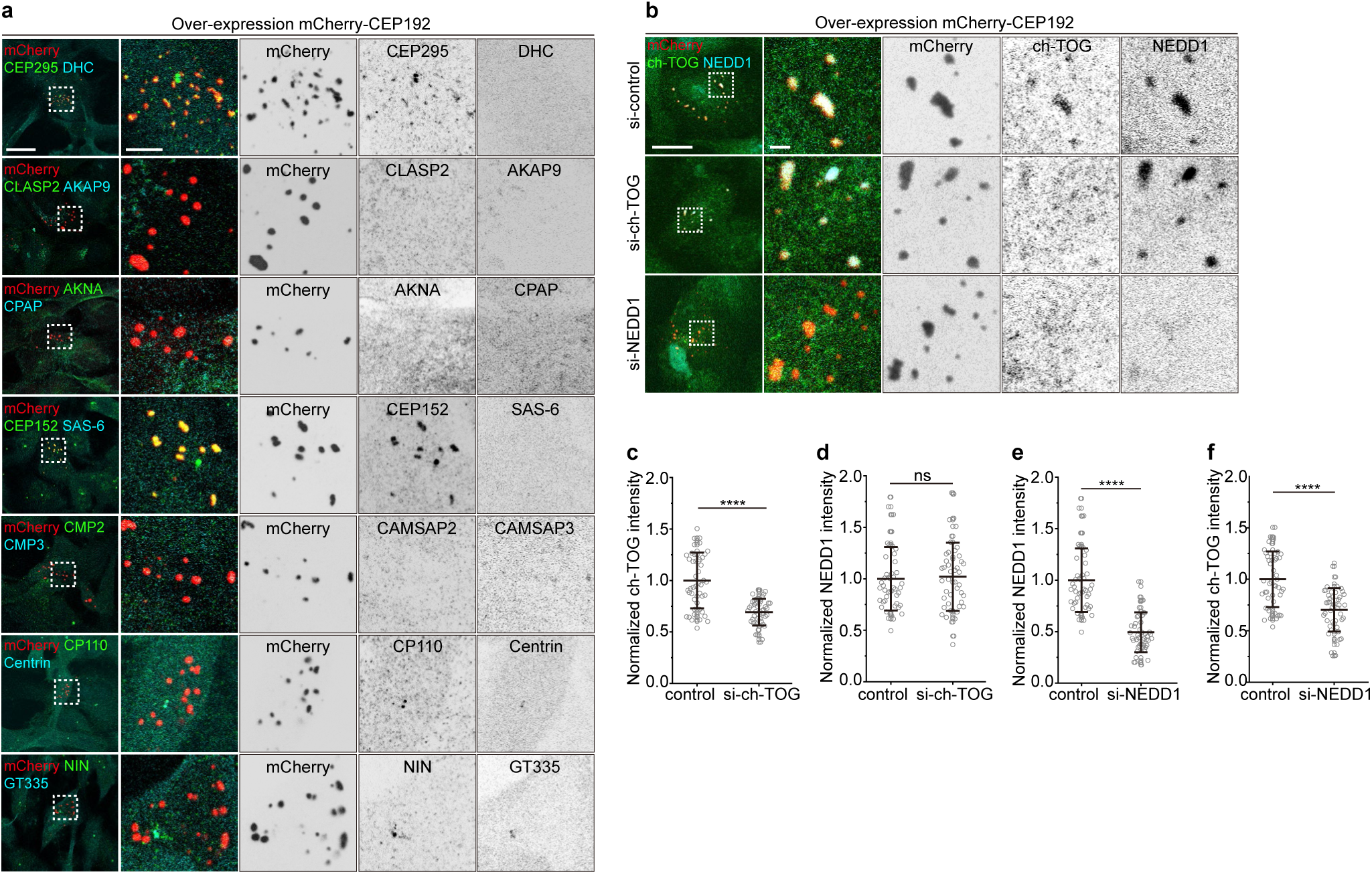
Components of CEP192 condensates in CEP192-overexpressing cells. **a,** Representative immunofluorescence images of mCherry-CEP192 condensates in wild type RPE1 cells stained for indicated proteins. Enlargements show merged and single channels of the boxed regions. Scale bar: 20 µm (main image); 5 µm (enlargement). **b,** Representative immunofluorescence images of mCherry-CEP192 condensates in wild type RPE1 cells. Cells were depleted indicated proteins and stained for mCherry (red), ch-TOG (green), and NEDD1 (cyan). Enlargements show merged and single channels of the boxed regions. Scale bar: 20 µm (main image); 5 µm (enlargement). **c, d,** Quantification of mean intensity of ch-TOG (mean ± s.d., **c**) and NEDD1 (mean ± s.d., **d**) in mCherry-CEP192 condensates depleted of ch-TOG, from the experiments shown in **b**. Values are normalized to the control average. The numbers of condensates analyzed (n) for each condition from three independent experiments are: for ch-TOG (**c**): control (n = 50, 15 cells), si-ch-TOG (n = 50, 20 cells); for NEDD1 (**d**): control (n = 50, 15 cells), si-ch-TOG (n = 49, 20 cells). Statistical significance was determined using one-way ANOVA with Tukey’s post hoc test for multiple comparisons (\*\*\*\**P* < 0.0001; ns, not significant, *P* = 0.7597). **e, f,** Quantification of mean intensity of NEDD1 (mean ± s.d., **e**) and ch-TOG (mean ± s.d., **f**) in mCherry-CEP192 condensates depleted of NEDD1, from the experiments shown in **b**. Values are normalized to the control average. The numbers of condensates analyzed (n) for each condition from three independent experiments are: for NEDD1 (**e**): control (n = 50, 15 cells), si-NEDD1 (n = 50, 16 cells); for ch-TOG (**f**): control (n = 50, 15 cells), si-NEDD1 (n = 50, 16 cells). Statistical significance was determined using one-way ANOVA with Tukey’s post hoc test for multiple comparisons (\*\*\*\**P* < 0.0001).

**Figure S5.**
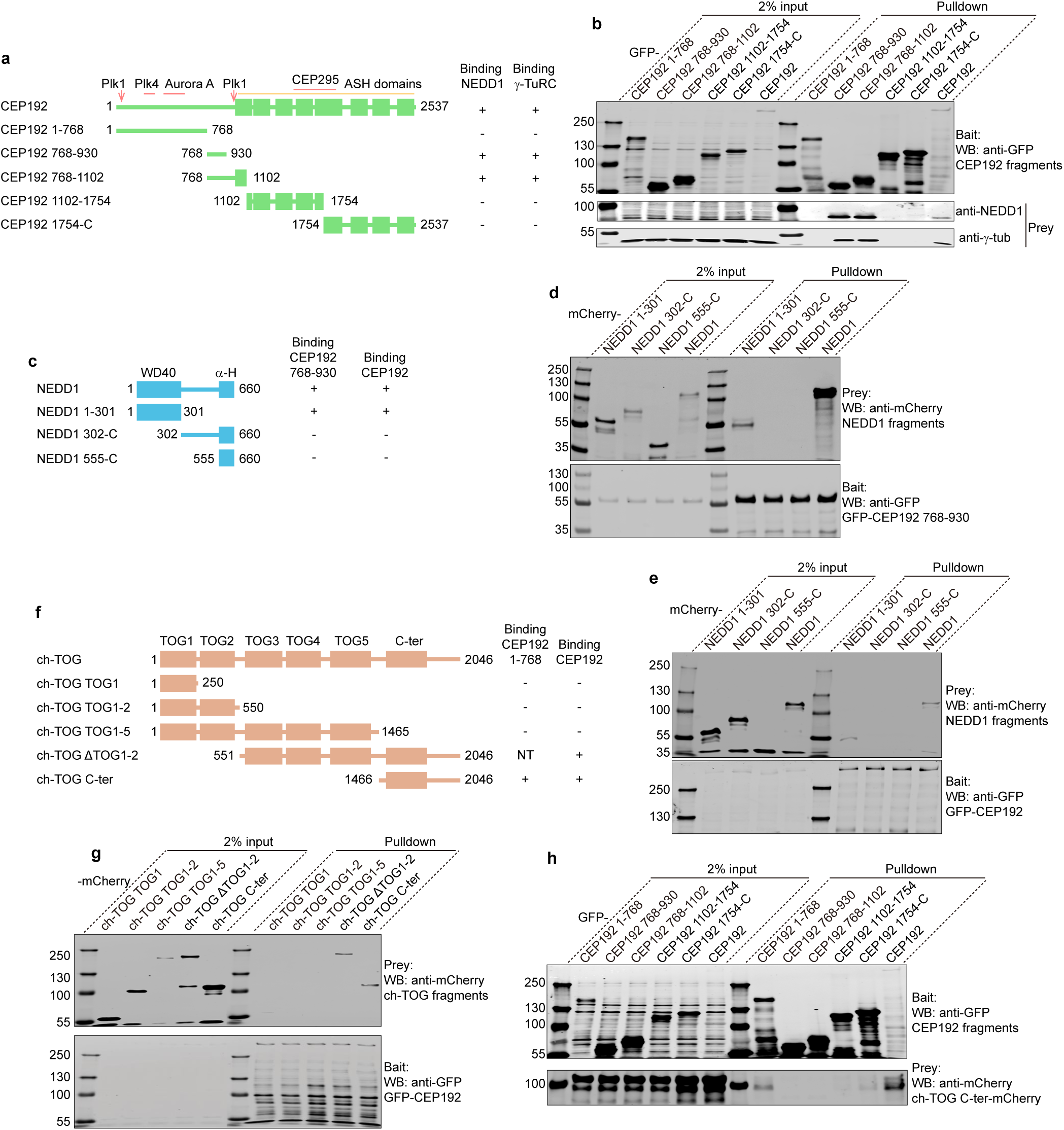
Mapping of the interactions between CEP192, NEDD1 and ch-TOG. **a, c, f,** Schemes of the domain organization of CEP192, NEDD1 and ch-TOG and various truncations used for pull-down assays. +, positive interaction; − no interaction detected, NT, not tested **b, d, e, g, h,** GFP pull-down assays with the indicated GFP constructs as bait and endogenous NEDD1 and γ-tubulin (**b**), mCherry-NEDD1 truncations (**d,e**) or ch-TOG-mCherry or its truncations as prey (**g,h**). Assays were performed using extracts of HEK293T cells expressing the indicated constructs and analyzed by Western blotting with antibodies against NEDD1, γ-tubulin, GFP and mCherry.

**Figure S6.**
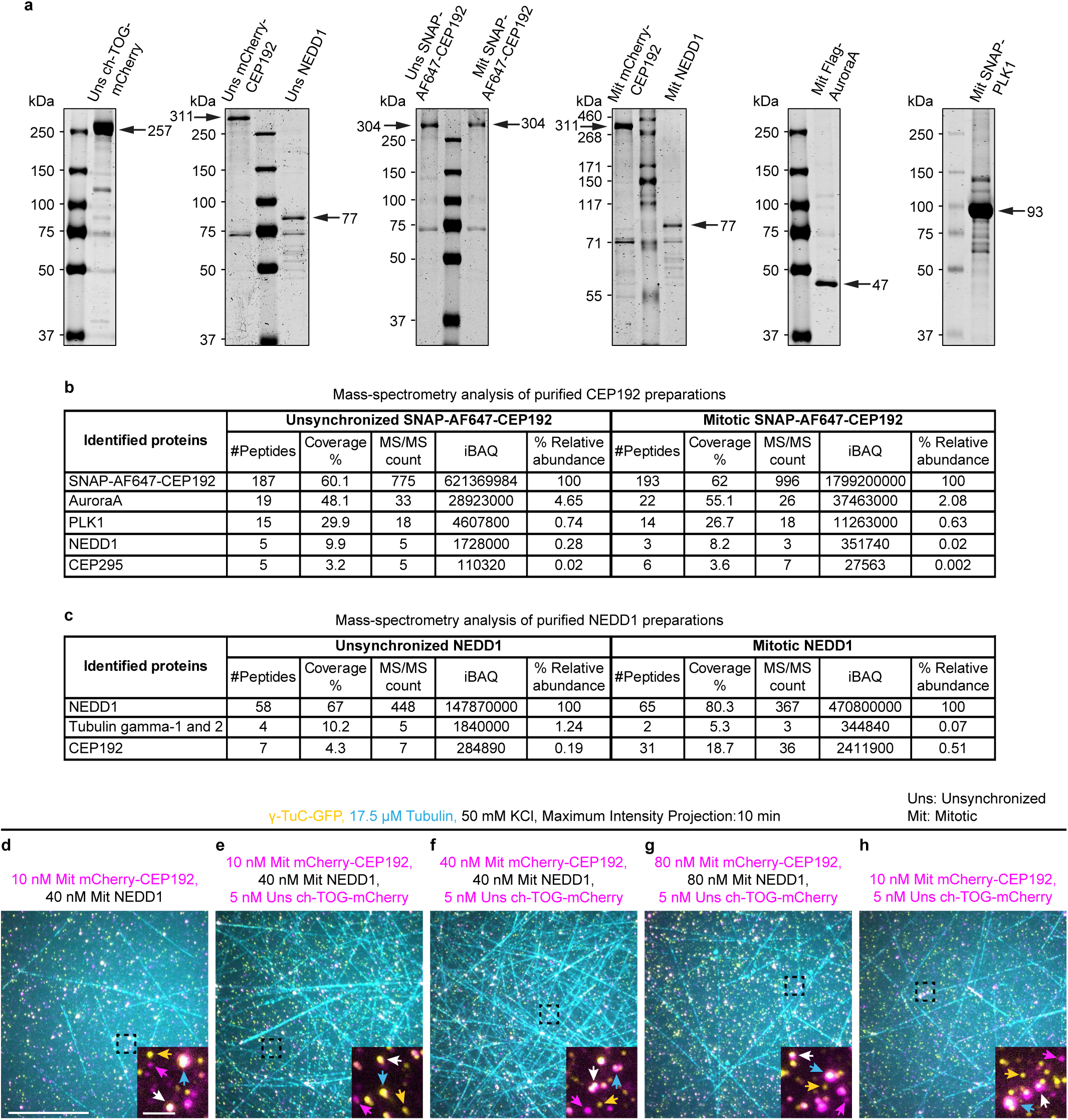
Characterization of purified recombinant proteins and the effective concentration of CEP192-NEDD1 complex. **a,** Purified Uns ch-TOG-mCherry-SII, Uns SII-mCherry-CEP192, Uns SII-NEDD1, Uns SII-SNAP-AF647-CEP192, Mit SII-SNAP-AF647-CEP192, Mit SII-mCherry-CEP192, Mit SII-NEDD1, Mit Flag-Aurora A and Mit SII-SNAP-PLK1 analyzed by Coomassie-stained SDS-PAGE. Black arrows indicate the purified recombinant protein bands at expected positions, and the numbers next to the arrows indicate their respective calculated molecular weight in kDa. **b,c,** List of proteins identified by mass spectrometry that were co-purified with Uns SII-SNAP-AF647-CEP192 (left) or Mit SII-SNAP-AF647-CEP192 (right) in (**b**) and Uns SII-NEDD1 (left) or Mit SII-NEDD1 (right) in **(c)**. Also, see **Table S2-5** for the full list of contaminants, including any potential interactors, which were not present in the top hits (< 0.5% relative abundance). **d-h**, Maximum intensity projections of 10 min videos, acquired after 10 min of incubation, showing MTs (cyan) nucleated from γ-TuCs (yellow, GCP3-GFP) under the indicated conditions: 10 nM Mit mCherry-CEP192 (magenta) together with 40 nM Mit NEDD1 (**d**); 10 nM (**e**) or 40 nM (**f**) Mit mCherry-CEP192 (magenta) together with 40 nM Mit NEDD1 and 5 nM Uns ch-TOG-mCherry (magenta); 80 nM Mit mCherry-CEP192 (magenta) together with 80 nM Mit NEDD1 and 5 nM Uns ch-TOG-mCherry (magenta) (**g**); 10 nM Mit mCherry-CEP192 (magenta) together with 5 nM Uns ch-TOG-mCherry (magenta) (**h**). Magnification is the same in **d-h**. Insets are the magnified view of the black squares showing colocalization of γ-TuC shown in yellow with CEP192 and ch-TOG shown here in magenta. Yellow arrowheads in the insets indicate non-colocalizing γ-TuCs; cyan arrowheads indicate active γ-TuCs; magenta arrowheads indicate non-colocalizing CEP192 puncta or the combined signal of CEP192 and ch-TOG puncta as CEP192 and ch-TOG were both mCherry-tagged proteins in these reconstitutions; white arrowheads indicate colocalizing γ-TuCs with CEP192 or CEP192-ch-TOG. Magnification is the same in the insets in **d-h.** In all the assay conditions, 2 mM ATP was added in the premixed γ-TuC solution before immobilization, as well as in the nucleation reaction mix with or without proteins. Scale bar: 20 µm, scale bar inset, 2 µm. Uns, proteins prepared from unsynchronized cells; Mit, proteins prepared from mitotic cells.

## Supplemental table legends

**Table S1.**
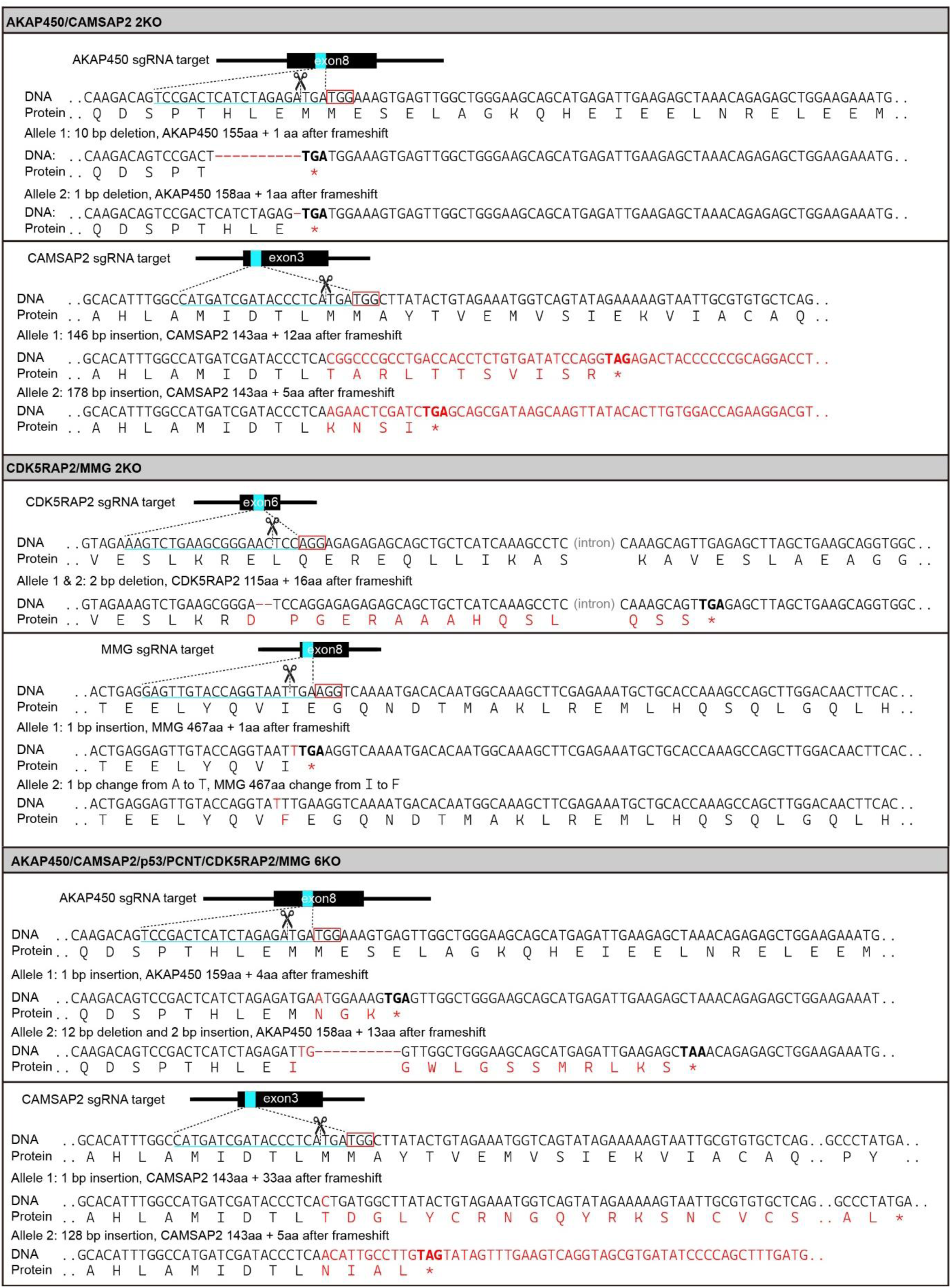
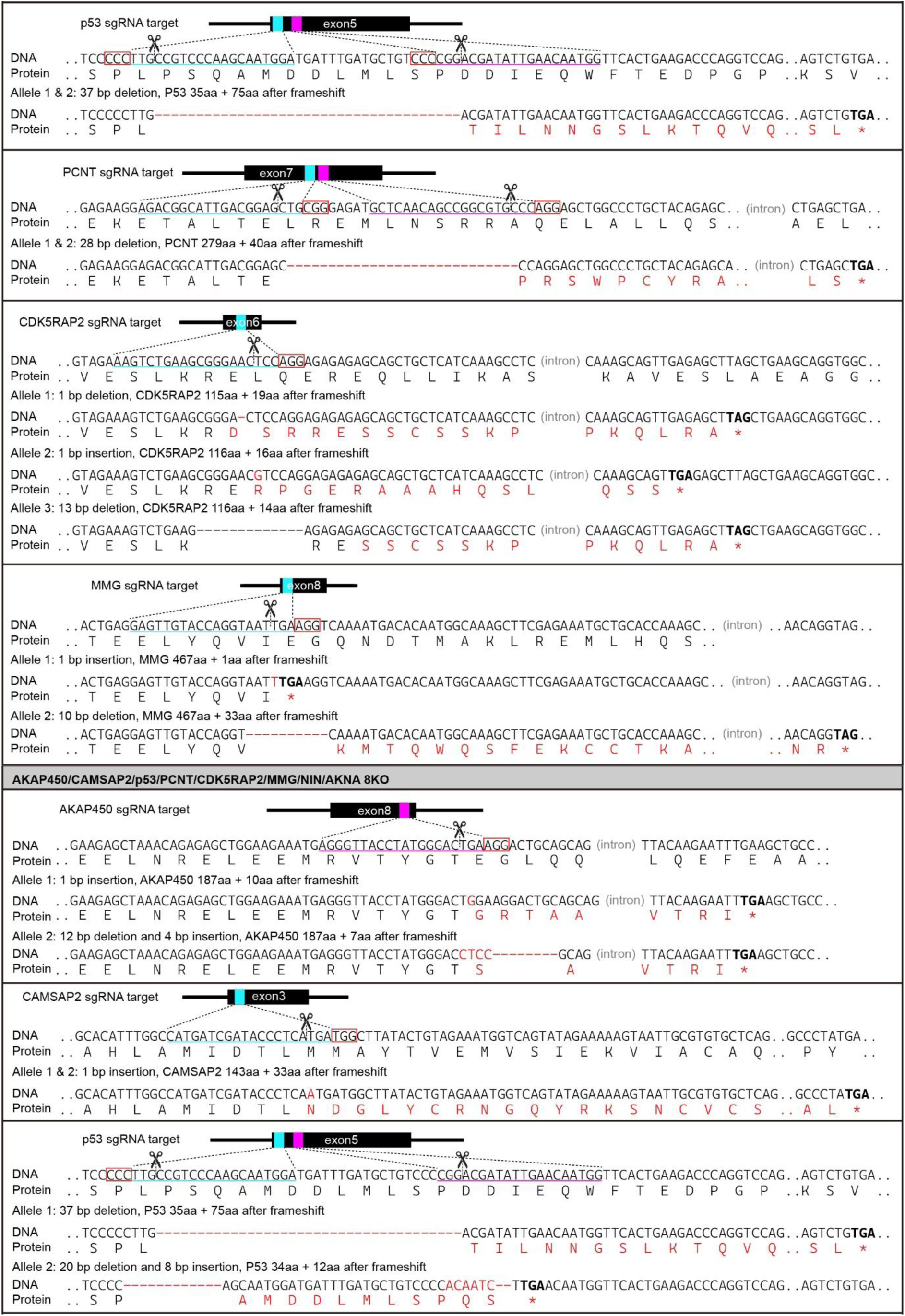
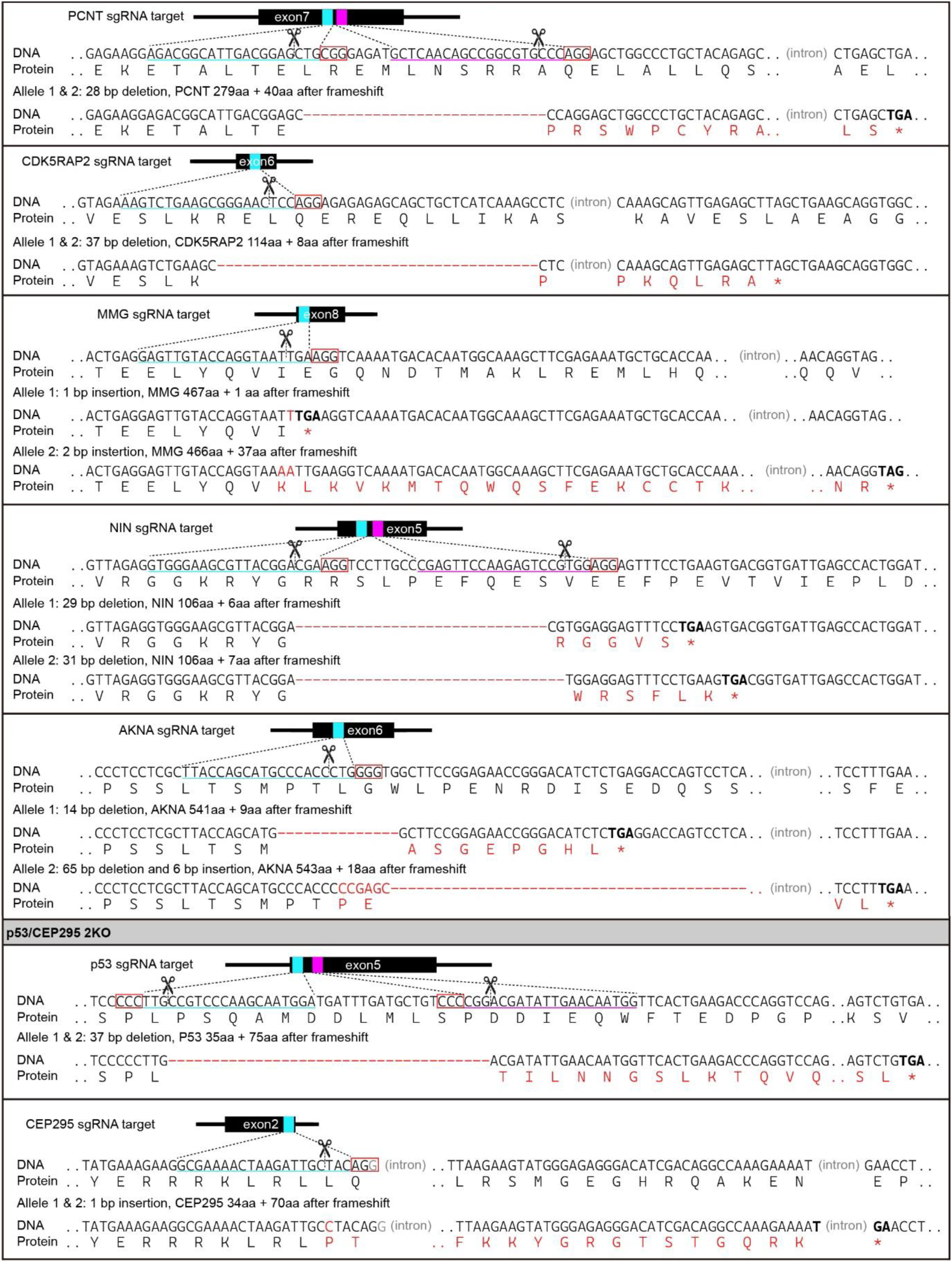
Gene sequencing of knockout RPE1 cell lines.

**Table S2. Mass-spectrometry analysis of SII-SNAP-AF647-CEP192, purified from unsynchronized cells**. List of proteins that were co-purified with SII-SNAP-AF647-CEP192 from unsynchronized cells.

**Table S3. Mass-spectrometry analysis of SII-NEDD1, purified from unsynchronized cells**. List of proteins that were co-purified with SII-NEDD1 from unsynchronized cells.

**Table S4. Mass-spectrometry analysis of SII-SNAP-AF647-CEP192, purified from mitotic cells**. List of proteins that were co-purified with SII-SNAP-AF647-CEP192 from mitotic cells.

**Table S5. Mass-spectrometry analysis of SII-NEDD1, purified from mitotic cells**. List of proteins that were co-purified with SII-NEDD1 from mitotic cells.

**Table S6.**
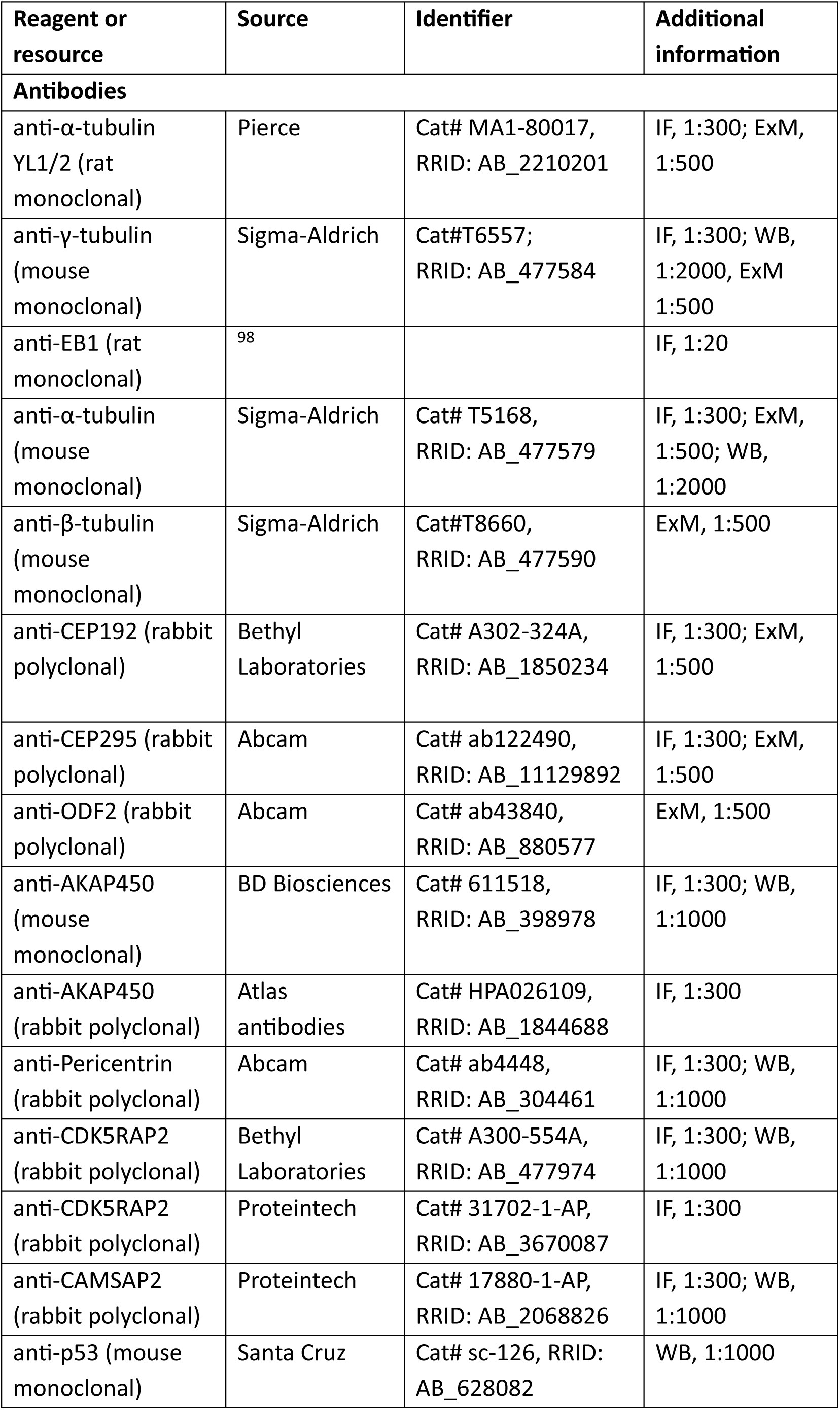
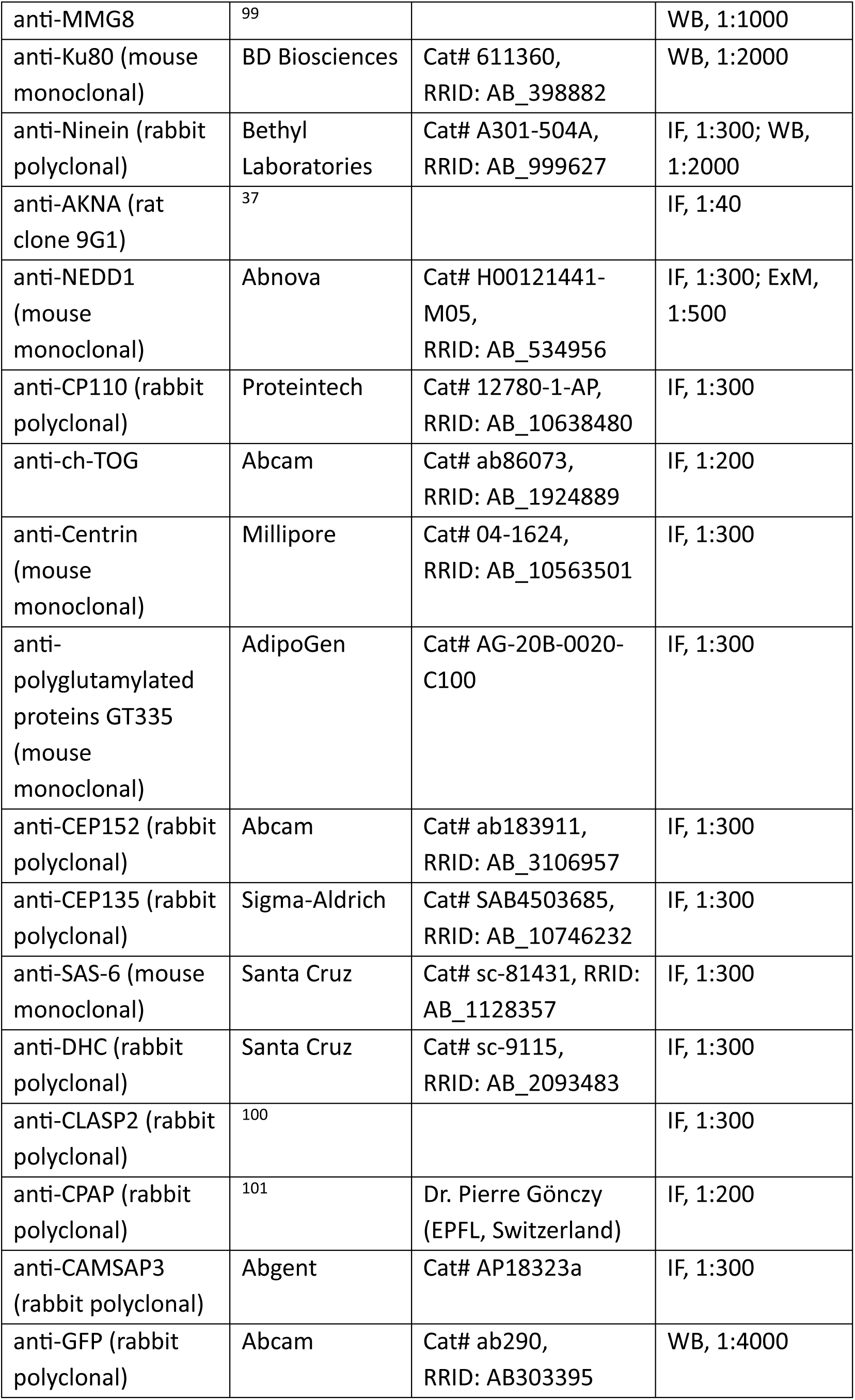
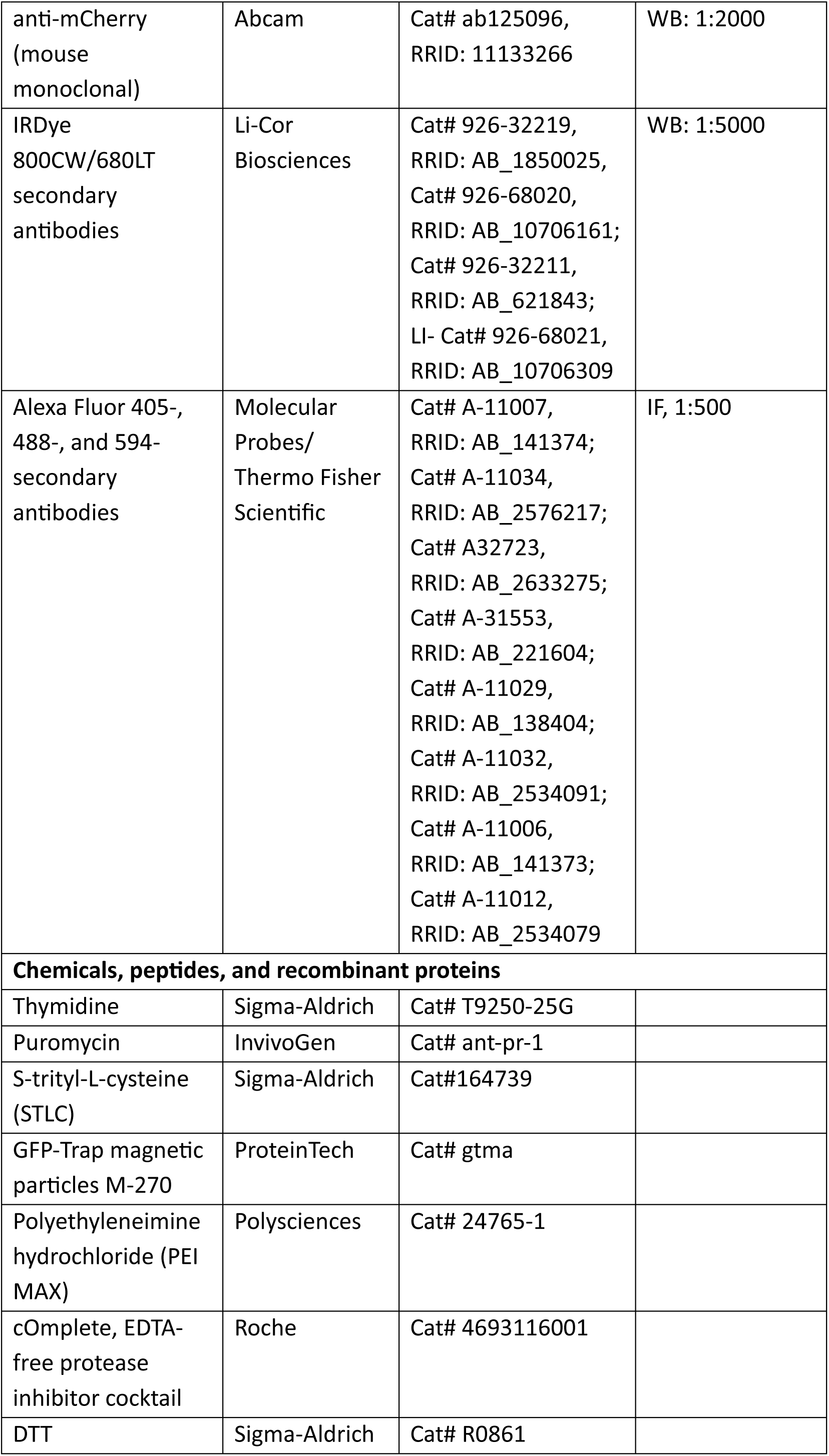
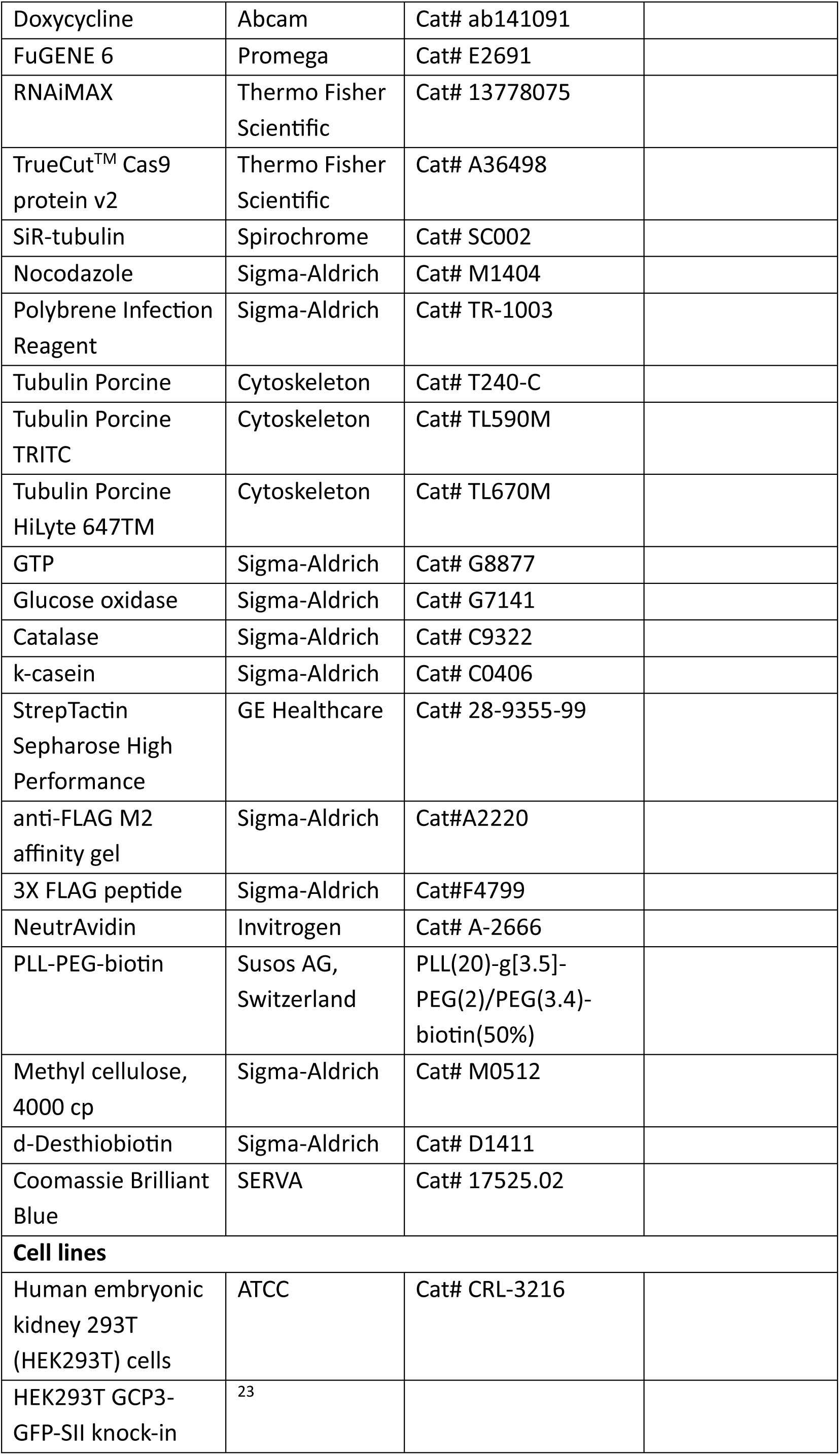
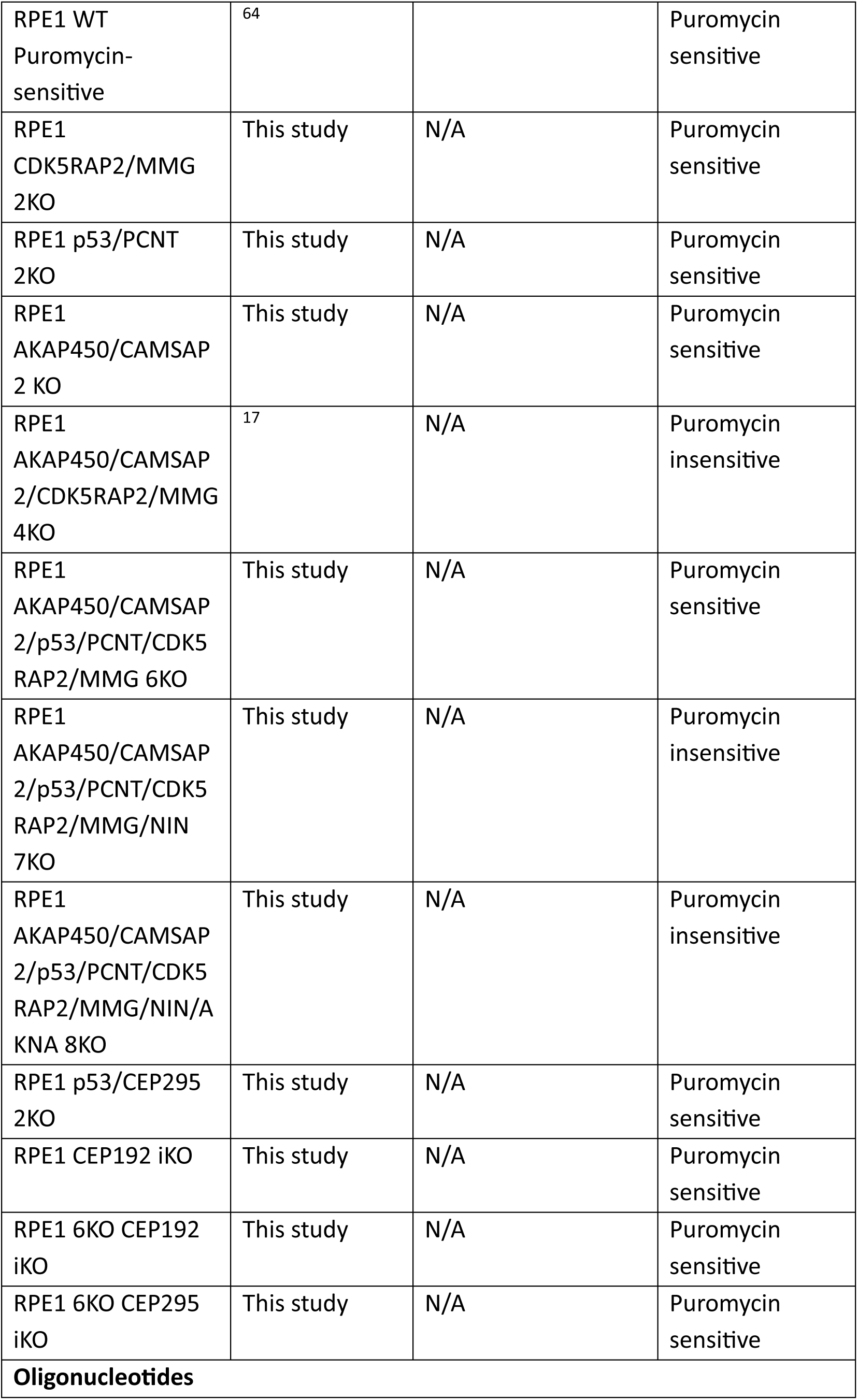
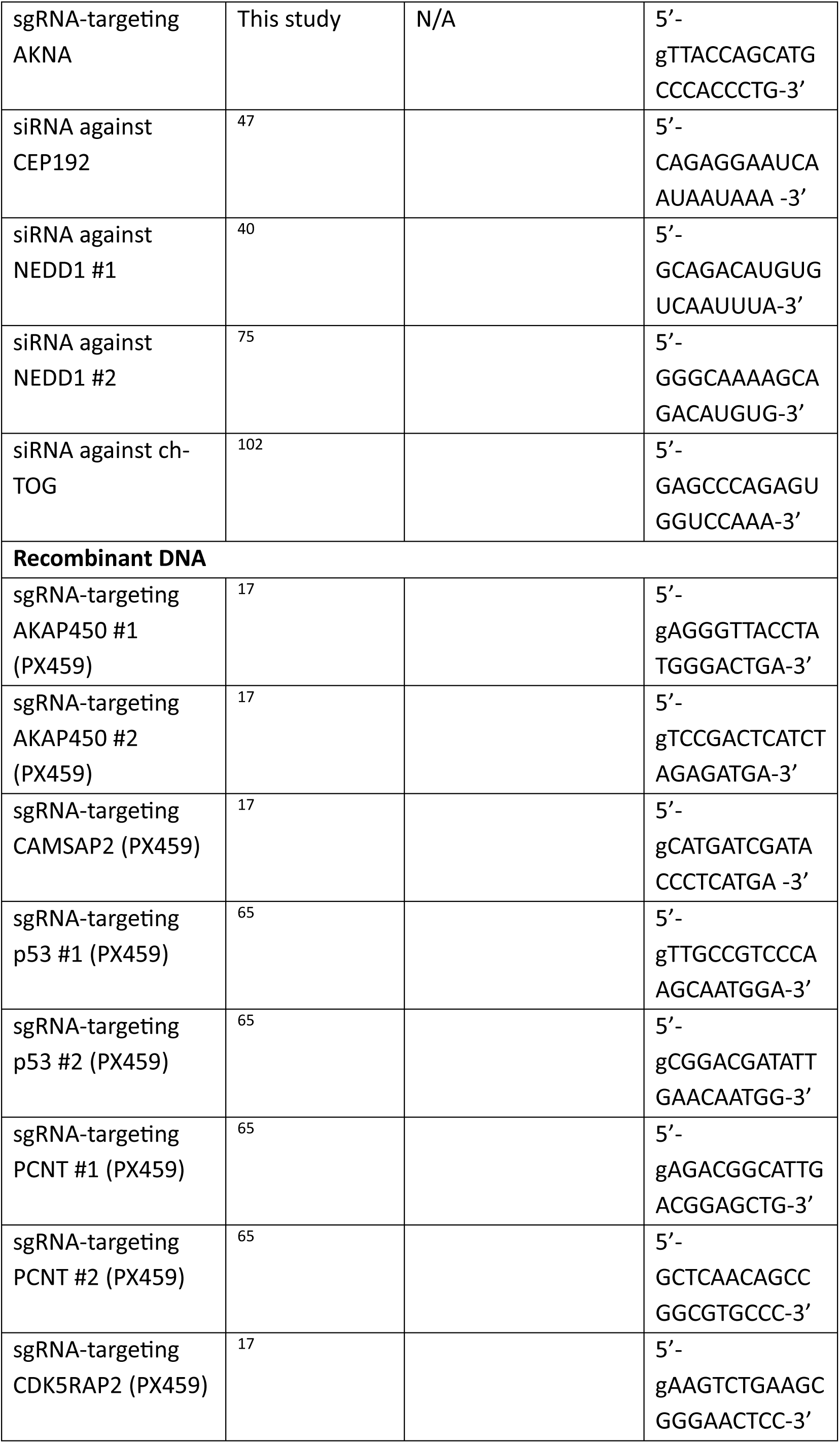
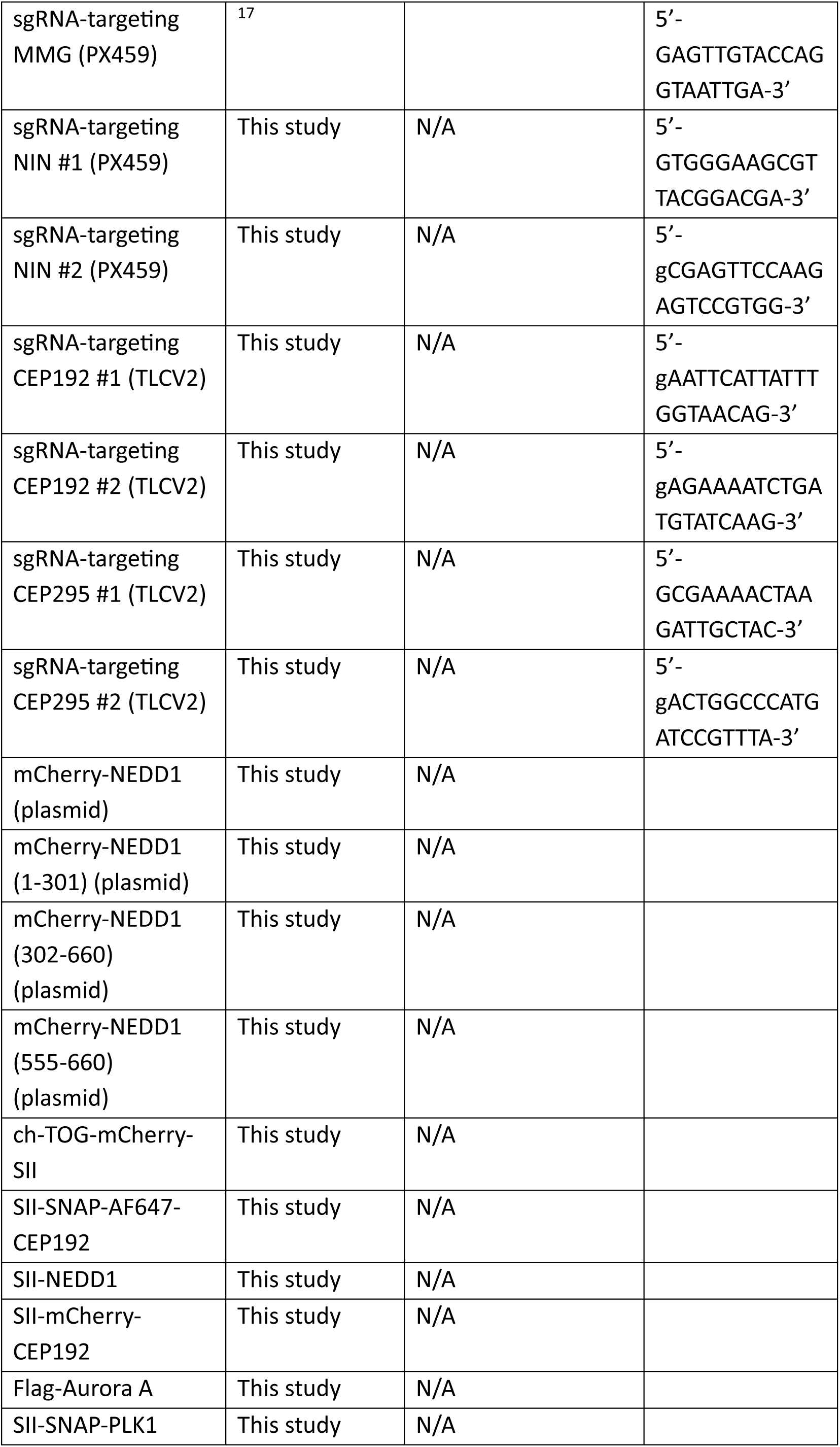
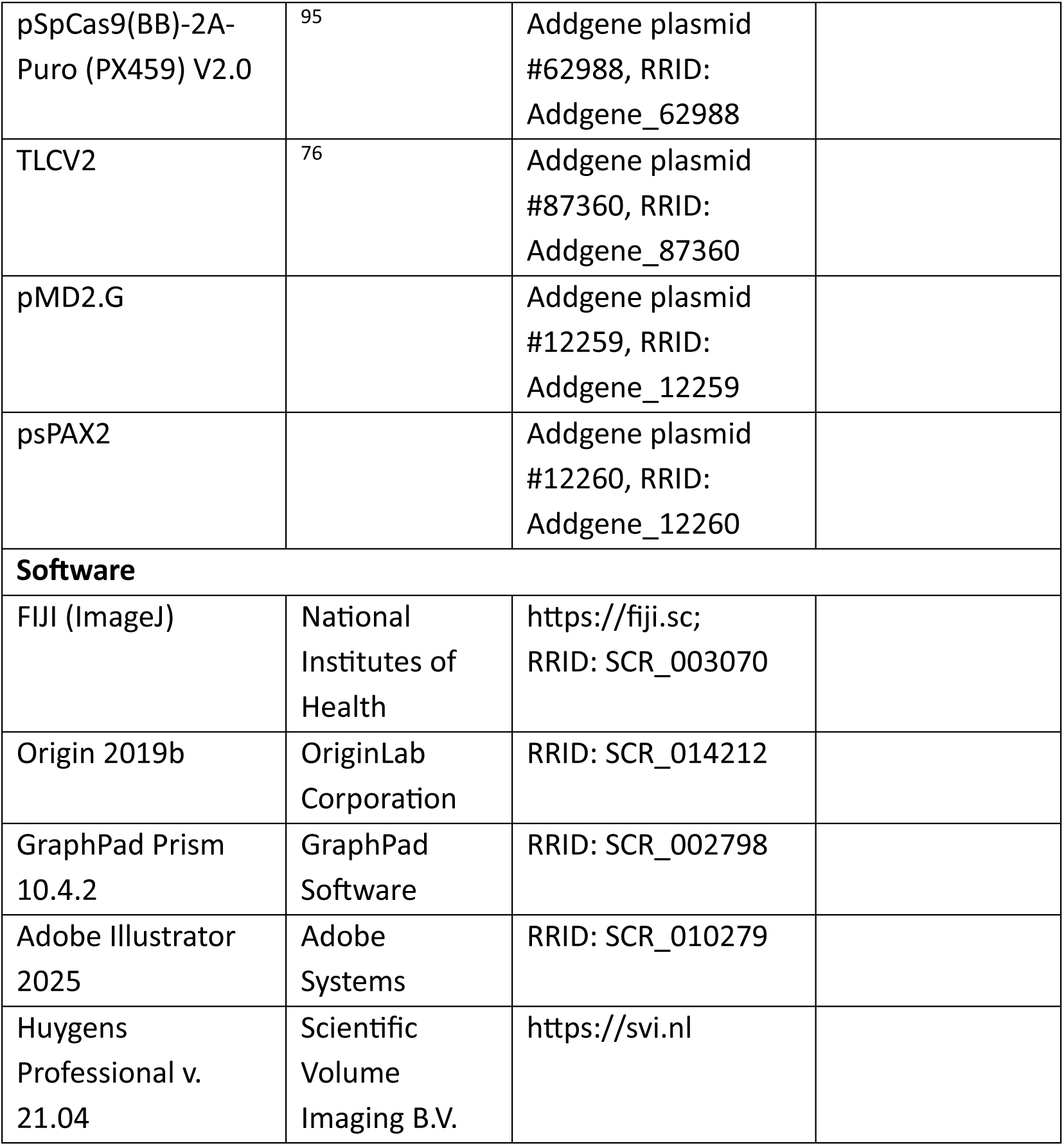
Key resources table.

## Source data

**Source data Fig. S1.**
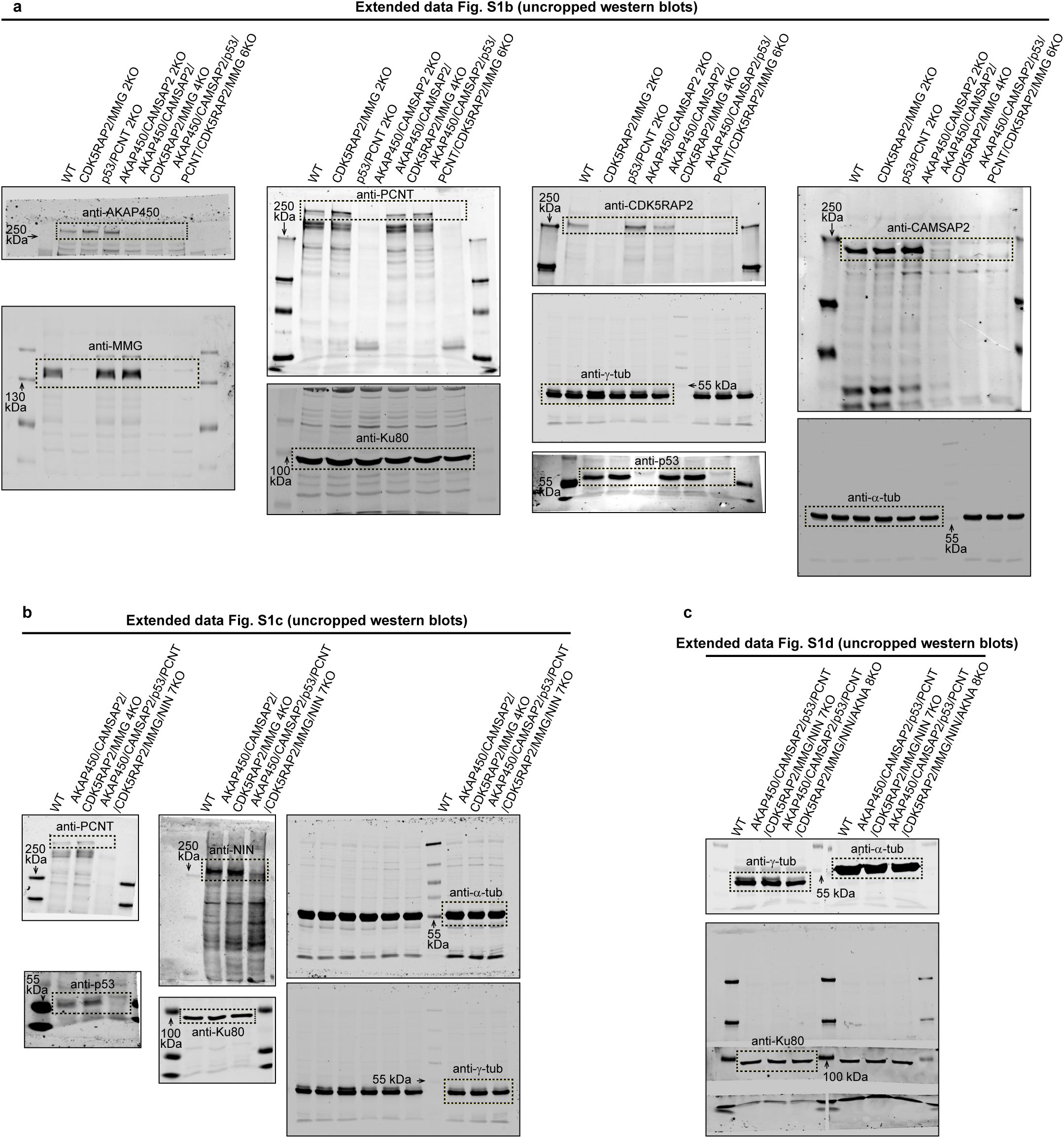
Unprocessed gels.

**Source data Fig. S5.**
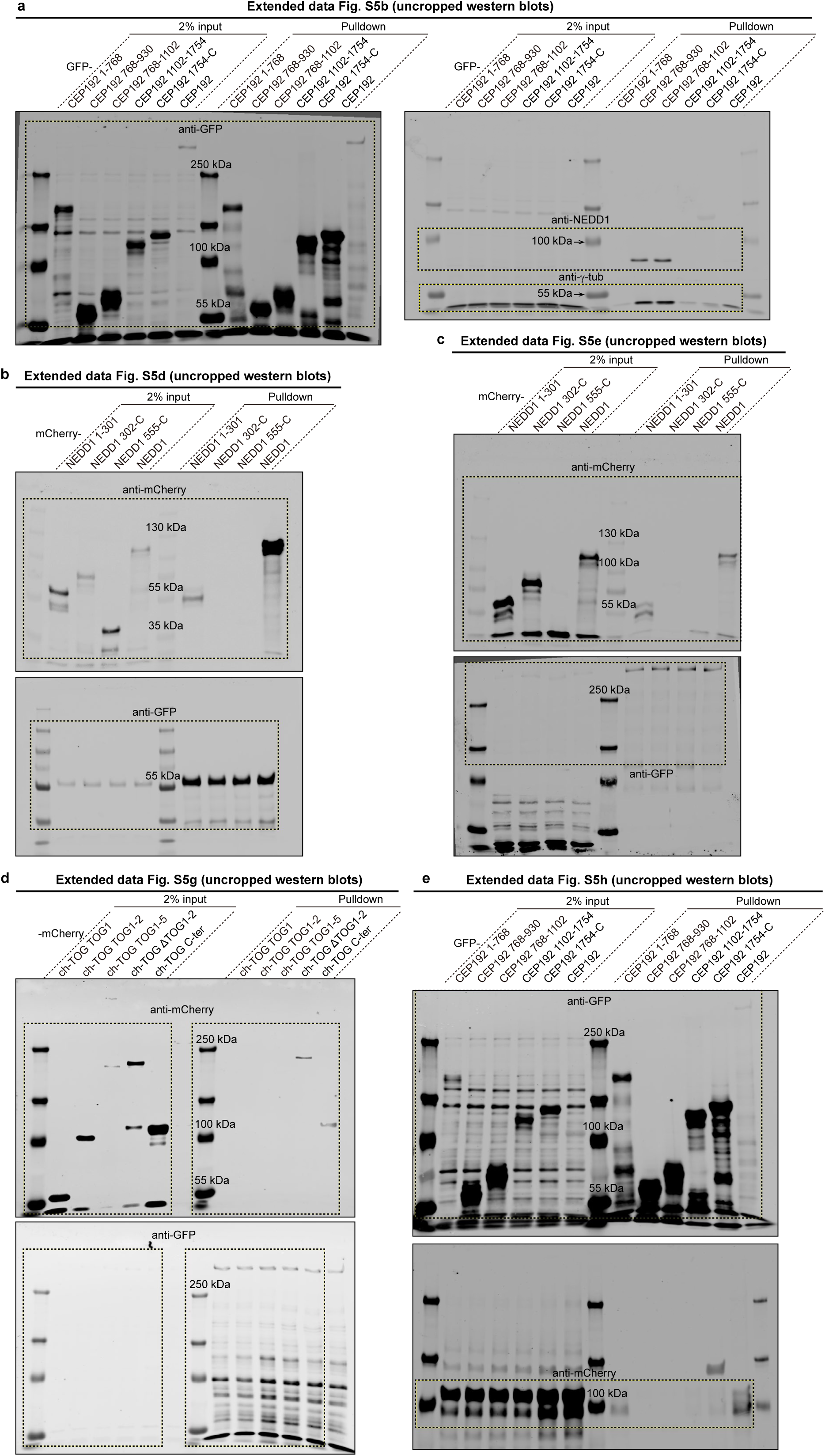
Unprocessed gels.

**Source data Fig. S6.**
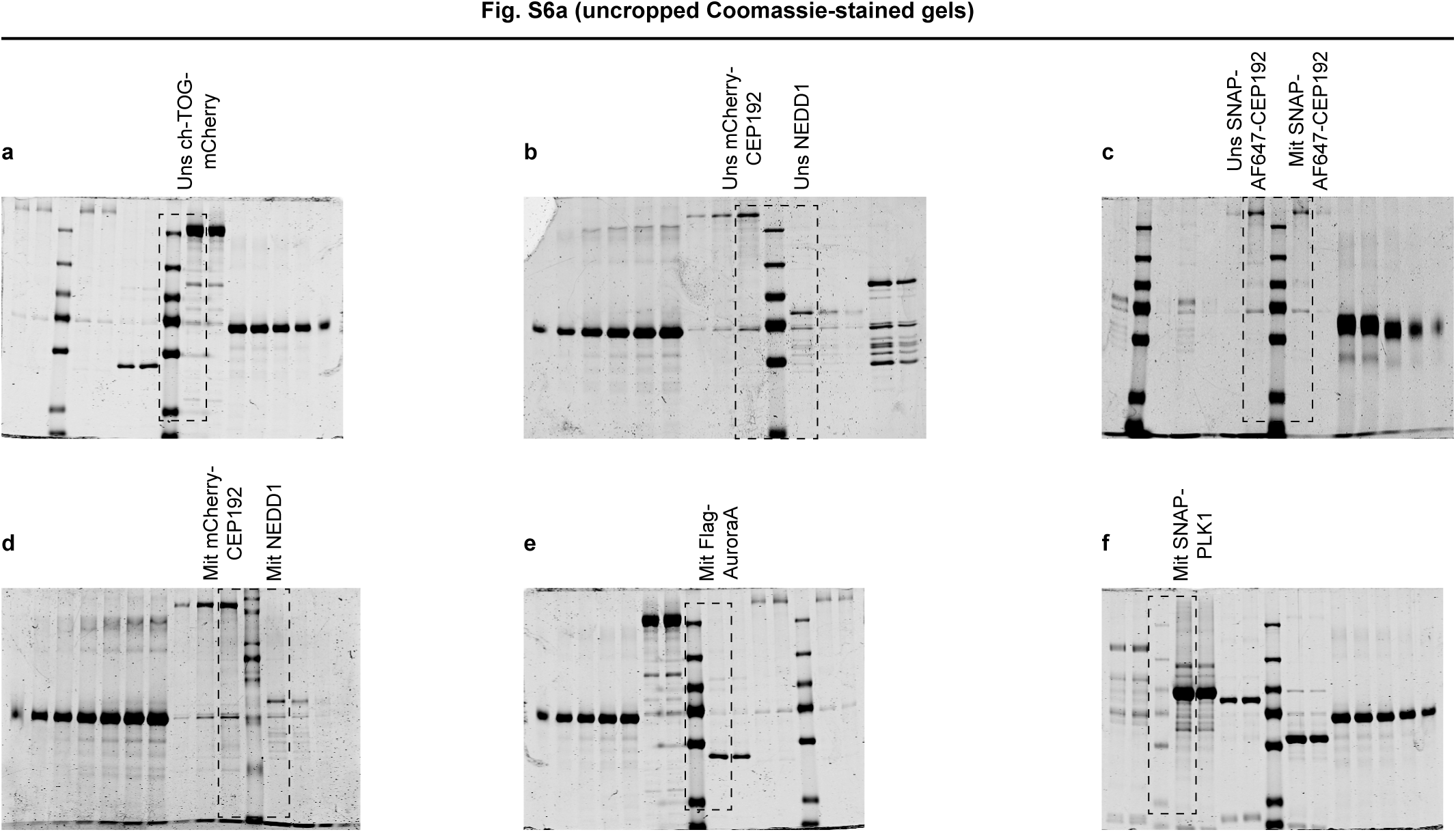
Unprocessed gels.

**Statistical source data.** Numerical and statistical source data.

## Methods

### DNA constructs

The constructs carrying full-length CEP192 (NM_032142.4, a gift from Andrew Holland, Johns Hopkins University School of Medicine), ch-TOG (NM_001008938.4) ^94^ and NEDD1 (NM_152905.4) ^85^ were confirmed by sequencing. Their truncations were cloned by either Gibson Assembly or T4 ligase-based ligation. To generate lentiviral vectors for inducible knockout of CEP192 and CEP295, we used the vector TLCV2 (Addgene Plasmid #87360) ^76^. This construct contains a constitutive sgRNA expression cassette, an inducible Cas9-T2A-GFP cassette controlled by Tet-On system, and a constitutive PuroR-P2-ArTetR cassette (Fig. S3a). The sgRNA sequences used for inducible knockouts in this study were as follows: CEP192, 5’-gAATTCATTATTTGGTAACAG-3’ and 5’-gAGAAAATCTGATGTATCAAG-3’; CEP295, 5’-GCGAAAACTAAGATTGCTAC-3’ and 5’-gACTGGCCCATGATCCGTTTA-3’.

The pSpCas9(BB)-2A-Puro (PX459, Addgene Plasmid # 62988) vector used for the CRISPR/Cas9 knockouts was acquired from Addgene ^95^ and used to clone the following 20-nucleotide targeting sequences: AKAP450, 5’-gAGGGTTACCTATGGGACTGA-3’, or 5’ gTCCGACTCATCTAGAGATGA-3’; CAMSAP2, 5’-gCATGATCGATACCCTCATGA-3’; P53, 5’-gTTGCCGTCCCAAGCAATGGA-3’ and 5’-gCGGACGATATTGAACAATGG-3’; PCNT, 5’-gAGACGGCATTGACGGAGCTG-3’ and 5’-GCTCAACAGCCGGCGTGCCC-3’; CDK5RAP2, 5’-gAAGTCTGAAGCGGGAACTCC-3’; MMG, 5’-GAGTTGTACCAGGTAATTGA-3’; NIN, 5’-GTGGGAAGCGTTACGGACGA-3’ and 5’-gCGAGTTCCAAGAGTCCGTGG-3’; AKNA, 5’-gTTACCAGCATGCCCACCCTG-3’.

For recombinant protein purification, we used previously described ch-TOG-mCherry-Twin-StrepII (SII) construct ^23^. Twin-StrepII-SNAP-CEP192, Twin-StrepII-mCherry-CEP192 and Twin-StrepII-NEDD1 were generated by cloning the full length constructs described above into modified pEGFP-C1 vectors with either a Twin-StrepII-SNAP, or Twin-StrepII-mCherry, or Twin-StrepII tag at the N-terminus. Flag-Aurora A in pcDNA3 plasmid was a gift from S. Lens (University Medical Center, Utrecht, The Netherlands). pRcCMV myc-Plk1 wt (Nigg RG6) construct (Addgene plasmid #41160) ^96^ was a gift from G. Kops (Hubrecht Institute and Oncode Institute, Utrecht, The Netherlands). Twin-StrepII-SNAP-PLK1 was made by cloning PLK1 into a modified pTT5 expression vector (Addgene plasmid #44006) with an N-terminal Twin-StrepII-SNAP tag.

### Cell lines, cell culture and transfection

We used previously published cell lines: hTERT RPE-1 (RPE1), HEK293T and HEK293T GCP3-GFP-Twin-StrepII (GCP3-GFP-SII) knock-in cells ^23^. Puromycin-sensitive RPE1 cells ^64^ were a gift from Andrew Holland (Johns Hopkins University School of Medicine). These cell lines were not found in the commonly misidentified cell lines database (International Cell Line Authentication Committee), and no further cell line authentication was performed. Cells were routinely checked for mycoplasma contamination using LT07-518 Mycoalert assay. RPE1 cells were cultured in Dulbecco’s Modified Eagle’s Medium DMEM/Ham’s F10 media (1:1) and HEK293T cells in DMEM alone, in both cases supplemented with 10% fetal bovine serum (FBS, GE Healthcare) and 1% penicillin/streptomycin (Sigma-Aldrich). All cells were cultured in tissue culture polystyrene flasks (Corning) and maintained in a humidified incubator at 37 °C with 5% CO_2_.

FuGENE 6 (Promega) was used to transfect RPE1 cells for CRISPR/Cas9 knockout generation and mCherry-CEP192 over-expression; RNAiMAX (Thermo Fisher Scientific) was used to transfect RPE1 cells with siRNAs; Polyethylenimine “Max” (PEI MAX, Polysciences) was used to transfect HEK293T cells for lentivirus packaging, pull-down experiments, and recombinant protein purifications. Transfections were performed according to the manufacturer’s instructions, using reagent/DNA or reagent/siRNA ratio within the recommended ranges. PEI MAX was used to transfect HEK293T cells with plasmids at 3:1 ratio of PEI MAX: plasmid for StrepTactin- and Flag-affinity-based protein purification.

### Drugs and drug treatments

The following drugs were used in this study: thymidine (Sigma-Aldrich), puromycin (InvivoGen), nocodazole (Sigma-Aldrich), and doxycycline (Abcam). Thymidine (2 mM) was used to arrest cells in interphase. For MT disassembly, cells were treated with 10 µM nocodazole for 1 hour on ice to achieve complete disassembly of stable MT fragments. Doxycycline (1 µg/ml) was used to switch on Cas9 expression in the Tet-On inducible knockout system.

### Generation of CRISPR/Cas9 knockout cell lines

RPE1 cells were transfected with the PX459 vectors containing the appropriate targeting sequences using FuGENE 6. One day after transfection, cells were subjected to selection with 15 µg/ml puromycin for up to 3 days. After selection, cells were allowed to recover in normal medium for approximately 7 days, and knockout efficiency was assessed by immunofluorescence staining. Depending on the observed efficiency, 50-500 individual clones were isolated and confirmed by immunofluorescence staining. The resulting single colonies were further characterized by Western blotting, immunostaining and genome sequencing.

Generation of AKNA knockout RPE1 cells was performed by electroporation of RPE1 cells with a Cas9/sgRNA mixture. In brief, 1 µl of Cas9 (20 µM, Thermo Fisher Scientific), 3.3 µl of sgRNA (30 µM, EditCo Bio) against AKNA, and 50 µl of Neurobasal medium were premixed on ice. A total of 1.5 × 10⁵ RPE1 cells were resuspended in 50 µl of Neurobasal medium and then added to the Cas9/sgRNA mixture to make a total volume of 100 µl. Cells were electroporated twice with a 20 ms, 1350 V pulse using an Invitrogen™ Neon™ Transfection System with 100 µl tips. The electroporated cells were plated in a 6-well plate and cultured for 1 day in DMEM/Ham’s F-10 medium (1:1) supplemented with 10% FBS, in the absence of penicillin/streptomycin.

### siRNA-mediated depletion

RPE1 cells seeded in 6-well plates were transfected with siRNAs using RNAiMAX. One day post-transfection, cells were allowed to recover in normal medium for 2 days. Cells were then detached from the 6-well plate and reseeded onto coverslips in 24-well plates. Cells in the 24-well plates were transfected with siRNAs again, and old medium was replaced with fresh medium one day post-transfection. After 2 days of culture, thymidine was added to arrest cells in interphase. Cells were harvested one day after thymidine blocking and subjected to immunofluorescence analysis.

### Lentivirus packaging

Lentiviruses were produced by PEI MAX-mediated co-transfection of HEK293T cells with the transfer vector, psPAX2 (packaging), and pMD2.G (envelope) (psPAX2 and pMD2.G were a gift from D. Trono (EPFL, Switzerland)). Viral supernatant was harvested 2-3 days post-transfection, filtered (0.45 µm), and precipitated overnight at 4 °C with PEG-6000. After centrifugation (1,500xg, 30 min), the pellet was resuspended in phosphate-buffered saline (PBS), aliquoted, and stored at −80 °C.

### Generation and analysis of inducible knockout cells

To generate inducible knockout of CEP192 and CEP295, two different lentiviruses each carrying distinct sgRNAs were packaged separately and then used together to infect parental RPE1 WT and 6KO cells. RPE1 cells were infected with the lentiviruses and incubated in complete medium supplemented with 8 µg/mL polybrene (Sigma-Aldrich). After 1 day, the medium was replaced with fresh complete medium. Starting 3 days post-transduction, cells were subjected to selection with puromycin at 25 µg/mL for up to 3 days, until most untransduced control cells (treated with the same concentration of antibiotic) were dead.

Inducible knockout cells were seeded onto coverslips in 24-well plates and treated with doxycycline for 3 days. Subsequently, thymidine was added to arrest cells in interphase. Cells were harvested one day after thymidine block and subjected to immunofluorescence analysis.

### MT disassembly and regrowth assay

Cells were treated with 10 µM nocodazole and incubated on ice for 1 hour to completely depolymerize MTs. Nocodazole was then removed by washing cells at least six times on ice with ice-cold complete medium. Subsequently, plates were transferred to a 37 °C water bath, and pre-warmed medium was added to each well to allow MT regrowth.

### Immunofluorescence cell staining and antibodies

Cells were fixed with pre-chilled (−20 °C) methanol for 5 min and permeabilized with 0.1% Triton X-100 in PBS for 2 min. Subsequent washing and labeling steps were carried out in PBS supplemented with 2% bovine serum albumin (BSA) and 0.05% Tween-20. Finally, slides were rinsed sequentially with 70% and 100% ethanol, air-dried, and mounted in ProLong™ Glass Antifade Mountant (Thermo Fisher Scientific).

### Ultrastructure Expansion Microscopy (U-ExM)

U-ExM was carried out as described by ^71^. Cells were incubated on ice for 60 min and pre-extracted for 90 s using 0.1% Triton X-100 in MRB80 (80 mM PIPES pH 6.8, 1 mM EGTA, and 4 mM MgCl₂). Cells were then washed three times with cold PBS and fixed for 5 hours with 1.4% paraformaldehyde (PFA) and 2.0% acrylamide in PBS at 37 °C. After a brief wash with PBS, a gelation solution (23% wt/vol sodium acrylate, 10% wt/vol acrylamide, 0.1% N,N′-methylenebisacrylamide, 1x PBS, 0.5% tetramethylethylenediamine (TEMED), and ammonium persulfate (APS)) was added to a rubber gelation chamber placed on a paraffin film (Parafilm)-covered glass slide and sealed with the coverslip containing the cells. The gel was allowed to polymerize for 1 hour at 37 °C.

After a brief wash with PBS, denaturation buffer (0.2 M SDS, 0.2 M NaCl, and 50 mM Tris-HCl, pH 9.0) was added and incubated for 15 min at room temperature, followed by further disruption for 90 min at 95 °C. The gel was then expanded in Milli-Q water for 30 min before being shrunk back through multiple rounds of PBS washes.

For immunostaining, the gel was incubated overnight at 4 °C with primary antibodies diluted in 1% BSA and 0.1% Triton X-100. Gels were washed extensively: 30 min in 1% BSA with 0.1% Triton X-100 in 1x PBS, 30 min in 0.1% Triton X-100 in 1x PBS, and three 1-hour washes in 0.1% Triton X-100 in 1x PBS. Gels were then incubated overnight at 4 °C with secondary antibodies in 1% BSA and 0.1% Triton X-100, followed by the same washing procedure used for the primary antibody. The gels were then transferred to Milli-Q water for final expansion. Prior to imaging, the gels were trimmed and mounted onto a plasma-cleaned, poly-L-lysine-treated coverslip.

A complete list of primary and secondary antibodies, including their dilutions, is provided in Table S6.

### Pull-down assays, immunoprecipitation, and Western blotting

HEK293T cells were seeded in 10 cm dishes and grown to 70–80% confluency. The cells were co-transfected with equal amounts of bait (GFP-tagged) and prey plasmid DNA using PEI MAX. 1 day after transfection, the cells were washed twice with ice-cold PBS and harvested. Cell pellets were lysed on ice for 30 min in 200 µL of lysis buffer (50 mM HEPES pH 7.4, 100 mM NaCl, 1 mM DTT, 0.5% Triton X-100) supplemented with EDTA-free protease inhibitor cocktail (Roche). The lysates were clarified by centrifugation at 14,000xg for 20 min at 4°C. The resulting supernatants were diluted with 300 µL of dilution buffer (50 mM HEPES pH 7.4, 100 mM NaCl, 1 mM DTT) containing protease inhibitors, and ~2% of this diluted lysate was mixed with 4x Laemmli sample buffer to be loaded as input. ChromoTek GFP-Trap Magnetic M-270 beads (ProteinTech) were pre-washed three times with wash buffer (50 mM HEPES pH 7.4, 100 mM NaCl, 1 mM DTT, 0.5% Triton X-100) using a DynaMag-2 magnet (Invitrogen). The remaining diluted lysates were incubated with the pre-washed beads for 1 h at 4°C with continuous rotation to allow efficient capture of the GFP-tagged bait and its interacting prey. Following incubation, the beads were collected magnetically and washed three times with wash buffer. After the final wash, the beads were resuspended in 2x Laemmli sample buffer, boiled at 95°C for 10 min to elute bound proteins, and then placed on a magnet to separate the beads. All samples were then loaded onto an SDS-PAGE gel.

Following SDS-PAGE, the unstained gels were transferred onto nitrocellulose membranes using a wet transfer system at 40 V for 12 h at 4°C. The membranes were blocked for 30 min with 2% BSA in PBS containing 0.05% Tween-20 (PBST) at room temperature. The membranes were then incubated overnight at 4°C with primary antibodies (provided in Table S6) diluted in blocking buffer. After primary antibody incubation, the membranes were washed three times (30 min each) with PBST. The membranes were subsequently incubated with appropriate IRDye-conjugated secondary antibodies (provided in Table S6) for 1 h at room temperature, protected from light. Following three additional washes with PBST (30 min each), the signal was detected using an Odyssey CLx infrared imaging system (Li-Cor Biosciences).

For Western blotting of knock out cell lines, cells were harvested from 10 cm dishes at 90% confluence. Protein extracts were prepared using a lysis buffer containing 50 mM HEPES (pH 7.4), 150 mM NaCl, 1 mM DTT, 0.5% Triton X-100, supplemented with EDTA-free protease inhibitor cocktail (Roche). Samples were separated by SDS-PAGE on polyacrylamide gels and transferred onto 0.45 µm nitrocellulose membranes (Sigma-Aldrich). Blocking was performed in 2% BSA in PBS for 30 min at room temperature. Membranes were incubated with primary antibodies overnight at 4 °C, washed three times with PBS containing 0.05% Tween-20, subsequently incubated with secondary antibodies for 1 hour at room temperature, and washed again three times with PBS containing 0.05% Tween-20. A complete list of primary and secondary antibodies, including their dilutions, is provided in Table S6. Membranes were imaged on an Odyssey CLx infrared imaging system using Image Studio software (version 5.2.5; Li-Cor Biosciences).

### Imaging of fixed cells and image analysis

Immunofluorescence imaging of fixed cells was performed using a Leica TCS SP8 STED 3X microscope equipped with an HC PL APO 100x/1.4 oil STED white objective, with white laser excitation at 488, 577, and 633 nm and depletion at 775 nm using a pulsed laser, driven by LAS X software. An internal Leica HyD hybrid detector with a time gate of 1 ≤ tg ≤ 8 ns was used, and the depletion laser power was set to 90% of maximum power. Images were acquired in 2D STED mode with a vortex phase mask.

Some fixed RPE1 cells were imaged on a Nikon Eclipse Ni upright wide-field fluorescence microscope equipped with a Nikon DS-Qi2 camera, an Intensilight CHGFI pre-centered fiber illuminator (Nikon), and ET-DAPI and ET-EGFP filters (Chroma), controlled by Nikon NIS-Br software. Slides were imaged using either a Plan Apo Lambda 60x numerical aperture (NA) 1.4 oil objective or a Plan Apo Lambda 100x NA 1.45 oil objective (Nikon).

### Live-cell imaging

Live fluorescence imaging was performed using spinning-disk confocal microscopy on an inverted Nikon Eclipse Ti-E research microscope (Nikon) equipped with the Perfect Focus System (Nikon), a Nikon Plan Apo VC 40x NA 1.3 objective, and a spinning-disk confocal scanner unit (CSU-X1-A1, Yokogawa). The system was also equipped with an ASI motorized stage with piezo top plate (MS-2000-XYZ, ASI), a Photometrics Evolve 512 EMCCD camera (Photometrics), and was controlled by MetaMorph 7.8 software (Molecular Devices). Vortran Stradus lasers (405 nm, 100 mW; 488 nm, 150 mW; 642 nm, 165 mW) and a Cobolt Jive 561 nm laser (110 mW) were used as light sources. The system was equipped with ET-DAPI (49000), ET-GFP (49002), ET-mCherry (49008), and ET-Cy5 (49006) filter sets (Chroma). To maintain cells at 37 °C with 5% CO₂, a stage-top incubator (INUBG2E-ZILCS, Tokai Hit) was used.

Inducible knockout cells were plated in 35 mm gridded dishes (Ibidi) and treated with 1 µg/mL doxycycline, 100 nM SiR-tubulin (Spirochrome), and 10 µM Verapamil (Spirochrome) before imaging. Cells were imaged with a 5-min interval and 200 ms exposure for 24 hours at 10% laser power.

Phase-contrast live-cell imaging was performed on a Nikon Ti microscope equipped with a Perfect Focus System (Nikon), a super-high-pressure mercury lamp (C-SHG1, Nikon), a Plan Apo 20x NA 0.75 objective, a CoolSNAP HQ2 CCD camera (Photometrics), and a motorized stage with piezo top plate (MS-2000-XYZ, ASI), all controlled by Micro-Manager software. A stage-top incubator (Tokai Hit) was used to maintain 37 °C and 5% CO₂. Cells were plated on round 25 mm coverslips, mounted in Attofluor Cell Chambers (Thermo Fisher Scientific), and imaged with a 2-min interval for 24 hours.

### Image analysis of fixed cells

To analyze centrosomal MT intensity in RPE1 cells, circular regions of interest (ROIs) of 10 µm² were drawn around each centrosome to obtain the mean intensity. Background intensity was measured as the minimum value within the ROI. Centrosomal MT intensity was calculated by subtracting the background intensity from the mean intensity. To analyze EB1 intensity at centrosomes following nocodazole washout in RPE1 cells, circular ROIs of 5 µm² were drawn to select the areas occupied by EB1. The mean intensity and the minimum value (background) of the EB1 signal were obtained. Centrosomal EB1 intensity was calculated by subtracting the background intensity from the mean intensity. To calculate total MT intensity, ROIs were manually drawn around individual cells, and the mean intensity of the α-tubulin signal was measured. Background intensity was obtained from an area within the nucleus, which lacks MTs.

To calculate protein intensity at centrosomes in RPE1 cells, a line profile was drawn across the centrosomal signal. The resulting intensity profile was fitted with a Gaussian function. Protein intensity was calculated by subtracting the background value from the peak value of the fitted Gaussian curve. To analyze the number of regrowing MTs (EB1 signal) following nocodazole washout, the Particle Analysis plugin in ImageJ was used. ROIs were drawn around individual cells, a threshold was applied at T = 13, and the minimum particle size parameter set to 8 pixel units. γ-Tubulin intensity was measured at one centrosome per cell, selecting the centrosome with the higher γ-tubulin signal.

To calculate the nucleation activity of CEP192 condensates, the number of MTs growing from each condensate was divided by the mean γ-tubulin intensity and the condensate area.

To quantify the distribution of γ-tubulin, NEDD1, and CEP192 along the centriole from the proximal to the distal end in U-ExM images, the mean fluorescence intensity was measured within a rectangular ROI of 200 nm in width, positioned adjacent to the centriole wall, while excluding signals originating from the procentriole.

To quantify the intraluminal NEDD1 and γ-tubulin intensity in U-ExM images, the mean fluorescence intensity was measured within a rectangular ROI of 100 nm in width, positioned over the lumen of the centriole.

U-ExM images of centrioles were deconvolved using Huygens Professional v. 21.04.

### Purification of γ-TuCs from HEK293T GCP3-GFP-SII knock-in cells

Human γ-TuCs used in the in vitro reconstitution assays were purified as described previously using Twin-StrepII-tag and Strep-Tactin affinity purification method ^23^. In brief, homozygous HEK293T GCP3-GFP-SII knock-in cells, maintained in dark, were collected from confluent eight 15 cm dishes, resuspended, and lysed in the lysis buffer (50 mM HEPES, 150 mM NaCl, 0.5% Triton X-100, 1 mM MgCl_2_, 1 mM EGTA, 0.1 mM GTP and 1 mM DTT, pH 7.4) supplemented with EDTA-free protease inhibitor cocktail (Roche). Cell lysate was cleared by centrifugation at 21,000xg for 20 min at 4°C. The resulting supernatant was incubated with equilibrated Strep-Tactin Sepharose beads (GE Healthcare) for 45 min at 4 °C. After binding, the beads were washed three times with the wash buffer (50 mM HEPES, 150 mM NaCl and 0.1% Triton X-100, 1 mM MgCl_2_, 1 mM EGTA, 0.1 mM GTP and 1 mM DTT, pH 7.4). γ-TuCs were then eluted for 15 min at 4 °C using elution buffer (50 mM HEPES, 150 mM NaCl, 0.05% Triton X-100, 1 mM MgCl_2_, 1 mM EGTA, 0.1 mM GTP, 1 mM DTT and 2.5 mM d-Desthiobiotin, pH 7.4). Purified γ-TuCs were aliquoted immediately, snap-frozen in liquid nitrogen, and stored at −80 °C. Wherever possible, samples were protected from light by covering tubes with aluminum foil throughout the purification process. The purity and composition of purified γ-TuCs using this method have been analyzed and published previously ^23^.

### Purification of recombinant proteins from HEK293T cells for in vitro reconstitution assays

Human ch-TOG-mCherry-SII, SII-SNAP-AF647-CEP192, SII-mCherry-CEP192, SII-NEDD1 and SII-SNAP-PLK1 used in the in vitro reconstitution assays were purified using same SII-tag and Strep-Tactin affinity purification method as described above for γ-TuC, but with modified buffers and steps. Briefly, HEK293T cells were transfected with 50 μg of the respective constructs per 15 cm dish and the culture medium was refreshed next day following transfection. Unsynchronized cells were harvested 2 days post-transfection from four 15 cm dishes per construct. For mitotic preparations of the recombinant proteins, cells were synchronized in mitosis using a kinesin-5 inhibitor, S-trityl-L-cysteine (STLC, Sigma-Aldrich) at 5 μM final concentration. STLC was added in the culture medium when the media were refreshed next day, and cells were cultured for 20 hours before harvesting rounded cells arrested in mitosis. Cells were resuspended and lysed in lysis buffer (50 mM HEPES, 300 mM NaCl, 0.5% Triton X-100, 1 mM MgCl_2_ and 1 mM EGTA (plus 1 mM ATP for PLK1), pH 7.4) supplemented with EDTA-free protease inhibitor cocktail (Roche). Cleared lysates were incubated with equilibrated Strep-Tactin Sepharose beads. After binding, beads were first washed five times using high salt (1 M NaCl) containing wash buffer (50 mM HEPES, 0.1% Triton X-100, 1 mM MgCl_2_ and 1 mM EGTA (plus 1 mM ATP for PLK1), pH 7.4), followed by three washes with the wash buffer containing 300 mM NaCl. For SNAP-tag labeling of CEP192 with Alexa Fluor 647 dye, washed beads were incubated for 1 hour in labeling buffer (50 μM Alexa Fluor 647 dye in 50 mM HEPES, 150 mM NaCl and 0.1% Triton X-100, 1 mM MgCl_2_, 1 mM EGTA and 1 mM DTT, pH 7.4). Excess dye was removed by washing the beads five times with 300 mM NaCl wash buffer. Proteins were eluted using elution buffer containing 50 mM HEPES, 150 mM NaCl, 0.05% Triton X-100, 1 mM MgCl_2_, 1 mM EGTA, 1 mM DTT and 2.5 mM d-Desthiobiotin (plus 1 mM ATP for PLK1), pH 7.4. Human Flag-Aurora A from mitotic cells was purified using the same protocol as PLK1, except that anti-FLAG M2 affinity agarose gel (Sigma-Aldrich) for affinity capture, and Flag-Aurora A was eluted with 0.2 mg/ml 3X FLAG Peptide (Sigma-Aldrich) instead of d-Desthiobiotin in the elution buffer. Purified proteins were aliquoted immediately, snap-frozen in liquid nitrogen, and stored at −80 °C. Sample purity was assessed by Coomassie-stained SDS-PAGE and mass spectrometry.

### Mass spectrometry

To assess the quality of purified recombinant proteins, samples were digested using the single-pot, solid-phase-enhanced sample preparation method (SP3) ^97^. In brief, 5.2 μg unsynchronized SII-SNAP-AF647-CEP192, 6.6 μg unsynchronized SII-NEDD1, 4.6 μg mitotic SII-SNAP-AF647-CEP192 and 5.8 μg mitotic SII-NEDD1 samples were first reduced and alkylated by adding 10 mM Tris(2-carboxyethyl) phosphine (TCEP) and 40 mM chloroacetamide (CAA). Samples were incubated first at 95 °C for 10 min, then in the dark for 20 min at room temperature (RT). Both hydrophilic and hydrophobic Cytiva Sera-Mag™ Carboxylate-Modified Magnetic Beads were used in a 1:1 ratio, 75 μg in total. The beads were washed twice with Milli-Q water, followed by incubation in 75% ethanol on a Thermomixer at 1000 rpm for 20 min at RT. Tubes were placed on a magnetic rack to pellet the beads, and the supernatant was removed. The samples were washed 2x with 80% ethanol, followed by 1x wash with 100% acetonitrile (ACN). Then, the beads were digested in 200 μl of 100 mM ammonium bicarbonate (AMBIC). The samples were sonicated for two min in a water bath and digested overnight using the proteases Trypsin (Promega) and Lys-C, in a 1:25 and 1:75 protease to protein ratio, respectively, in a Thermomixer at 1000 rpm, at 37 °C. The samples were spun down and acidified in 5% trifluoracetic acid (TFA). The beads were immobilized on the magnetic rack, and the acidified peptide solutions were transferred to new Eppendorf tubes. Before LC-MS analysis, all digested peptide samples were eluted and dried in a vacuum centrifuge. Samples were analyzed by reversed phase nLC-MS/MS using an Ultimate 3000 UHPLC coupled to a Thermo Scientific^TM^ Orbitrap Exploris 480. Digested peptides were dissolved in 2% formic acid (FA) and separated on a 50-cm reversed-phase analytical column, with an integrated emitter, packed in-house (ReproSilPur C18-AQ 1.9 µm resin (Dr. Maisch GmbH), 75 µm inner diameter), using a linear gradient with buffer A (0.1% FA) and buffer B (80% acetonitrile, 0.1% FA) ranging from 4-44% B at a flow rate of 300 nL/min over 45 min. The Acclaim Pepmap 100 C18 (5 mm × 0.3 mm, 5 μm, ThermoFisher Scientific) trap column was operated at a temperature of 32 °C, and the analytical column at a temperature of 50 °C. The column was then washed with 99% Buffer B for 5 min, followed by a re-equilibration step with 4% Buffer B for 10 min.

For all samples, the MS data was acquired using a DDA method. MS1 scan parameters in profile mode were set to 60,000 resolution, a scan range of 375-1600 m/z, the automatic gain control (AGC) target to standard and an automatic maximum injection time. The MS2 scan parameters were set at 15,000 resolution, with a standard AGC target, an automatic maximum injection time, and an isolation window of 1.4 m/z. Scans were acquired with an initial fixed mass of 120 m/z, over a mass range of 200-2000 and a normalized collision energy (NCE) of 28. Precursor ions were selected for fragmentation using a 1 s acquisition cycle, a 10 s dynamic exclusion time and a charge filter between +2 to +6.

Raw data were processed in MaxQuant-Andromeda software (v.2.4.7.0) for protein identification, using the iBAQ values for quantification. The MS/MS spectra were searched against the UniProtKB Human Proteome database (organism_id: 9606, reviewed, canonical & isoform FASTA) together with a separate FASTA file containing the recombinant protein. Default parameters were applied for the precursor mass tolerance (20 p.p.m first search, 4.5 p.p.m. main search), and the False-Discovery Rate (FDR) was set at 1%. Up to three missed cleavages were allowed for each proteases. Carbamidomethylation of cysteine residues was set as a fixed modification, while methionine oxidation and protein N-terminal acetylation were set as variable modifications, with a maximum of five modifications allowed per peptide. The iBAQ values generated by MaxQuant were imported into the Perseus software (v.2.0.11.0), where protein groups labelled as ‘Reverse’ (false positives), ‘Contaminants’, and ‘Only identified by site’, as well as those identified by a single peptide, were filtered out. The iBAQ intensities of co-purified proteins within a sample were then normalized to the iBAQ intensity of the corresponding recombinant protein in that sample to calculate relative iBAQ ratios. This ratio provided an estimate of abundance of all the proteins present in the sample relative to the recombinant protein.

### In vitro reconstitution assays

#### In vitro reconstitution of MT nucleation with purified γ-TuCs

In vitro reconstitution assays were carried out as described previously ^23^. Flow chambers were prepared by attaching plasma-cleaned glass coverslips (18 x 18 mm coverslips) to microscopic slides using double-sided tape. Chambers were functionalized by incubation with 0.2 mg/ml PLL-PEG-biotin (Susos AG, Switzerland) for 5 min, followed by incubation with 1 mg/ml NeutrAvidin (Invitrogen) in MRB80 buffer (pH 6.8, 80 mM K-PIPES, 1 mM EGTA and 4 mM MgCl_2_) for another 5 min. Next, 25 nM biotinylated-anti-GFP nanobody was immobilized on the coverslip via biotin-NeutrAvidin links and incubated for 5 min. During this step, purified γ-TuC-GFP was diluted 10-fold in MRB80 buffer, with or without 2 mM ATP, and preincubated for 3 min with the indicated purified proteins (except ch-TOG) or with ATP-added MRB80 buffer as a control. The γ-TuC mix was then immobilized on the GFP-nanobody-coated coverslips by a 3 min incubation in the flow chamber. Unbound γ-TuCs were removed by washing with MRB80 buffer, and subsequently, the flow chambers were blocked with 0.8 mg/ml k-casein to prevent non-specific protein binding. MT nucleation mix with or without indicated proteins (MRB80 buffer supplemented with 17 μM unlabeled and 0.5 μM Rhodamine- or HiLyte647-labeled porcine brain tubulin, 50 mM KCl, 1 mM GTP, 0.5 mg/ml k-casein, 0.1% methylcellulose, oxygen scavenger mix (50 mM glucose, 400 μg/ml glucose-oxidase, 200 μg/ml catalase and 4 mM DTT), and with or without 2 mM ATP) were added to the flow chambers after centrifugation at 119,000xg for 5 min in an ultracentrifuge (Beckman Airfuge). Protein concentrations, determined by comparison to protein standard on Coomassie Blue-stained gel, and nucleation mix compositions are provided in the corresponding figure or figure legends. Chambers were sealed with high-vacuum silicone grease (Dow Corning) and equilibrated with the nucleation reactions for 2 min at 30 °C on Total Internal Reflection Fluorescence (TIRF) microscope stage. 10 min time-lapse videos after 10 min incubation with the nucleation mix were acquired. All tubulin products were from Cytoskeleton.

### Image acquisition, processing and data analysis of in vitro reconstitution experiments

#### TIRF microscopy

In vitro assays were imaged using an iLas2 TIRF microscopy system (Roper Scientific, Evry, France), a dual laser illuminator providing azimuthal spinning TIRF illumination. The system was mounted on a Nikon Eclipse Ti-E inverted microscope equipped with a perfect focus system, a Nikon Apo TIRF 100x 1.49 N.A. oil immersion objective, a CoolSNAP MYO M-USB-14-AC CCD camera (Roper Scientific), and an Evolve mono FW DELTA 512×512 EMCCD camera (Roper Scientific) fitted with a 2.5x intermediate lens (Nikon C mount adaptor). Excitation was provided by 150 mW 488 nm,100 mW 561 nm and 110 mW 642 nm lasers, using Chroma 49002 and 49008 filter sets. Image acquisition was controlled with MetaMorph 7.10.2.240 software (Molecular Devices). A stage top incubator (INUBG2E-ZILCS, Tokai Hit) maintained the temperature at 30 °C during imaging. 10 min time-lapse images were acquired with the CoolSNAP Myo CCD camera (Roper Scientific) at 5 s time intervals, using 300 ms exposure times for 561 nm and 642 nm illuminations and 500 ms for 488 nm excitation. The CoolSNAP Myo CCD camera provided a final pixel size of 0.045 μm.

#### Quantification of γ-TuC nucleation efficiency

For each independent nucleation assay, γ-TuCs that either nucleated MTs or were associated with a pre-existing MT during the 10 min observation of the time-lapse video were manually scored as active γ-TuCs. Total number of γ-TuC (GCP3-GFP) particles were detected and quantified from the first or second frame of that time-lapse video using the open-source ImageJ plugin ComDet v.0.5.6 (https://github.com/ekatrukha/ComDet). Nucleation efficiency was calculated as the percentage of active γ-TuCs relative to the total number of detected γ-TuCs.

#### Quantification of colocalization frequency of γ-TuC with CEP192 or ch-TOG

All the γ-TuC (GCP3-GFP) particles and CEP192 or ch-TOG particles immobilized on the coverslip within the field of view were detected from their respective channels from the first frame or second frame of the 10 min time-lapse movie using the ImageJ plugin ComDet mentioned above. The colocalization percentage between γ-TuC and CEP192 or ch-TOG was provided by the colocalization function of the ComDet plugin. In the parameters, the maximum distance between the colocalized spots was set to 4 pixels.

## Statistical analysis

All statistical details of experiments including the definitions, exact values of number of measurements, the number of independent experiments and measurements (or cells analyzed), precision measures and statistical tests performed are provided in the corresponding figure panels or legends. Data processing and statistical analysis were done in Excel, OriginPro 2019b 9.6.5.169 and GraphPad Prism 10.4.2. Significance was defined as: ns = not significant for *p* > 0.05, \**p* < 0.05, \*\**p* < 0.01, \*\*\**p* < 0.001 and \*\*\*\**p* < 0.0001.

## Data availability

Processed mass spectrometry data that support the findings of this study are provided as supplementary tables in Excel format. Source data are provided with this study. All data that support the conclusions are either available in the manuscript itself or available from the authors on request.

## Code availability

Fiji macros used in this study are available online at https://github.com/ekatrukha/ComDet.

